# Genetic dissection of glutamatergic neuron subpopulations and developmental trajectories in the cerebral cortex

**DOI:** 10.1101/2020.04.22.054064

**Authors:** Katherine S. Matho, Dhananjay Huilgol, William Galbavy, Miao He, Gukhan Kim, Xu An, Jiangteng Lu, Priscilla Wu, Daniela J. Di Bella, Ashwin S. Shetty, Ramesh Palaniswamy, Joshua Hatfield, Ricardo Raudales, Arun Narasimhan, Eric Gamache, Jesse Levine, Jason Tucciarone, Eric Szelenyi, Julie A. Harris, Partha P. Mitra, Pavel Osten, Paola Arlotta, Z. Josh Huang

## Abstract

Diverse types of glutamatergic pyramidal neurons (PyNs) mediate the myriad processing streams and output channels of the cerebral cortex, yet all derive from neural progenitors of the embryonic dorsal telencephalon. Here, we establish genetic strategies and tools for dissecting and fate mapping PyN subpopulations based on their developmental and molecular programs. We leverage key transcription factors and effector genes to systematically target the temporal patterning programs in progenitors and differentiation programs in postmitotic neurons. We generated over a dozen temporally inducible mouse Cre and Flp knock-in driver lines to enable combinatorial targeting of major progenitor types and projection classes. Intersectional converter lines confer viral access to specific subsets defined by developmental origin, marker expression, anatomical location and projection targets. These strategies establish an experimental framework for understanding the hierarchical organization and developmental trajectory of PyN subpopulations that assemble cortical processing networks and output channels.

## INTRODUCTION

Glutamatergic pyramidal neurons (PyNs) constitute the large majority (∼80% in rodents) of nerve cells in the cerebral cortex. In contrast to the mostly short-range projecting GABAergic inhibitory interneurons that form cortical local circuitry, long-range projecting PyNs mediate the myriad inter-areal processing streams and all of the cortical output channels^1–3^. Traditionally, the vast number of PyNs have been broadly classified into a handful of major classes according to their laminar location or broad axon projection patterns, such as intratelencephalic (IT) and extratelencephalic (ET or corticofugal), which further comprises subcerebral (including pyramidal tract-PT) and cortico-thalamic (CT) PyNs^1, 4^. It is increasingly evident that within these major subclasses, subsets of PyNs form specific local and long-range connectivity, linking discrete cortical microcircuits to separate cortical subnetworks and output channels^1, 5^. Recent single cell RNA sequencing (scRNAseq) analysis suggests over a hundred PyN transcriptomic types from just two cortical areas^6^. As a satisfying definition of PyN types requires correspondence across molecular, anatomical and physiological properties, reliable experimental access to specific PyN subpopulations is not only necessary for such multi-modal analysis but also for further exploring their circuit function. However, genetic tools for PyN subpopulations are severely limited, and the experimental strategy for a systematic dissection of PyN subpopulations that reflect their circuit organization has not been established.

Despite their immense number and diversity, all PyNs are generated from neural progenitors seeded in the embryonic dorsal telencephalon. Classic studies have established that a protomap of regionally differentiated radial glial progenitors (RGs) lining the cerebral ventricle undergo repeated rounds of asymmetric divisions and give rise to radial clones of PyNs, which are sequentially deployed to the cortex in a largely inside-out order^7^. RGs generate PyNs either directly or indirectly through intermediate progenitors (IPs), which divide symmetrically to generate pairs of PyNs^8^. Recent scRNAseq analysis at multiple developmental stages have revealed a set of key temporal patterning genes that drive lineage progression in RGs, and a largely conserved differentiation program that unfolds in successively generated postmitotic neurons^9^. Together, these studies delineate an outline of the cellular and molecular logic of PyN development^3, 9, 10^. However, the diversity and lineage organization of cortical progenitors remain unresolved and debated^11–14^, and it is unclear whether different RG subpopulations (e.g. marked by transcription factor expression) and neurogenetic mechanisms (direct *vs.* indirect) contribute, ultimately, to projection-defined PyN subpopulations. Resolving these fundamental issues requires genetic fate mapping tools with sufficient progenitor type resolution and temporal precision that enable tracking the developmental trajectory of PyNs.

Among the challenges of establishing genetic access to PyN types, the first and foremost is specificity (with an appropriate granularity) and the second is comprehensiveness. Although a large number of transgenic driver lines have been generated that label subsets of PyNs, only a handful are specific and reliable enough for neural circuit analysis^15, 16^. More importantly, as there is no simple relationship between single gene expression and “neuron types”, it is not clear whether and how a systematic dissection of PyN types can be achieved. Here, we have generated and characterized a set of 16 *Cre* and *Flp* gene knockin mouse driver lines for targeting PyN progenitors and subpopulations, guided by knowledge in their developmental genetic programs. Orthogonal and temporally inducible drivers enable combinatorial targeting of major progenitor types (e.g. RGs, IPs, neurogenic progenitors) and projection classes (e.g. CT, PT, IT subclasses and subpopulations within). Intersectional converter lines confer viral access to specific subsets defined by cell birth time, marker expression, anatomical location and projection targets. These tools and strategies establish an experimental framework for accessing hierarchically organized PyN projection types at progressively finer resolution. They will not only facilitate systematic multi-modal analysis of PyN subpopulations to define cortical circuit elements but also enable tracking their developmental trajectories toward elucidating the organization and assembly of the processing networks and output channels of the cerebral hemisphere, including the cortex, hippocampus, and basolateral amygdala.

## RESULTS

Our overarching strategy is to engage the precise spatiotemporal gene expression patterns of the specification and differentiation programs to target biologically relevant progenitor subsets, PyN subpopulations, and their developmental trajectories (Fig. 1). As such our approach differs from previous efforts^15, 16^ in several significant ways. First, we use gene knockin (KI) instead of transgenic approach to precisely recapitulate the endogenous pattern of transcription factors (TFs) and effector genes with well-characterized roles demonstrated by developmental genetic studies. Second, as temporal patterning of TF expression is an essential and evolutionarily conserved mechanism for generating cell type diversity, we generated mostly inducible *CreER* driver lines to leverage this versatile and efficient strategy. Third, the high reliability of the KI approach allows strategic design of combinatorial schemes across developmental stages. Fourth, intersectional reporter and converter lines integrate multiple developmental genetic properties and further engage anatomic features for viral targeting of highly specific PyN types.

**Figure 1:**
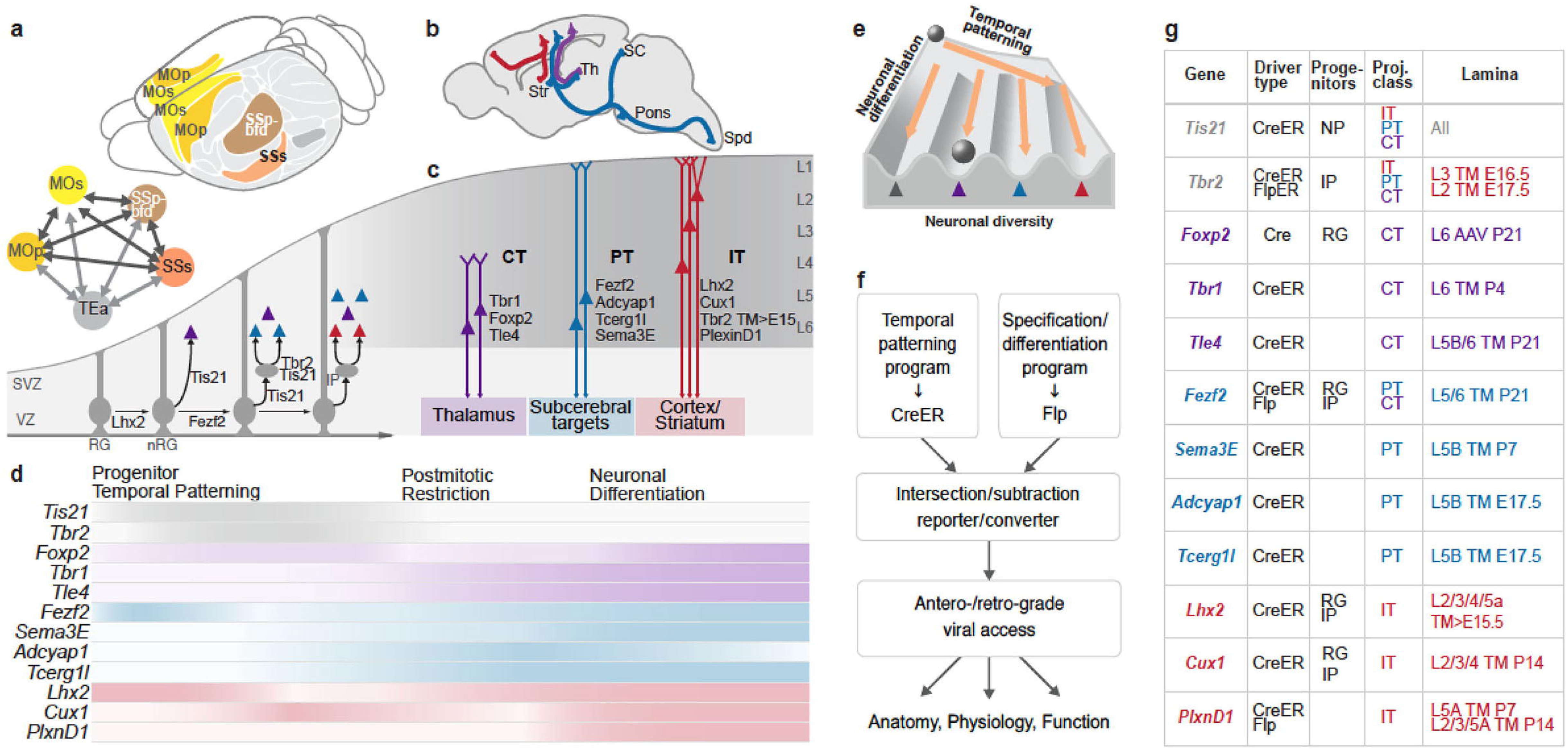
Strategies and tools for targeting PyN subpopulations and developmental trajectories. **a,** The mouse neocortex consists of multiple cortical areas (upper) that are preferentially interconnected into functional subnetworks, exemplified by the colored sensorimotor network (lower) comprising the primary and secondary sensory (SSp-bfd, SSs) and motor (MOp, MOs) and temporal association (TEa) cortex. **b,** A schematic sagittal brain section depicting several major PyN projection classes that mediate intratelencephalic streams (IT-red; cortical and striatal) and cortical output channels (PT-blue, CT-purple). Str, striatum; Th, thalamus; SC, superior colliculus; Spd, spinal cord. **c,** A schematic developmental trajectory of PyNs. During lineage progression, radial glial cells (RGs) undergo direct neurogenesis and indirect neurogenesis through intermediate progenitors (IPs) to produce all the laminar and projection types. Genes that are used to target progenitor types and PyN subpopulations are listed according to their cellular expression patterns. VZ, ventricular zone; SVZ, subventricular zone; nRG, neurogenic RG. **d,** Temporal expression patterns of genes used for generating driver lines across major stages of PyN development. Colors correspond to projection classes in b; intensity gradients represent approximate changes of expression levels across development for each gene. **e,** The specification and differentiation of PyNs along two cardinal axes: a temporal patterning program in dividing RGs that seeds the successive ground states of their progeny and a largely conserved differentiation program in sequentially born postmitotic PyNs that unfolds the diverse identities (modified with permission^9^). **f,** A strategy for combinatorial targeting of specific PyN projection types defined by developmental origin, marker expression, anatomical location and projection targets. This is achieved by converting the intersection of two gene expression patterns through Cre and Flp drivers into viral access that further engage anatomical properties. **g.** List of new driver lines generated and characterized in this study. NP, neurogenic progenitor.

All KI drivers (except as noted) are generated by in-frame insertion of *CreER* or *Flp* coding cassettes before the STOP codon of an endogenous gene, separated by a T2A self-cleaving peptide sequence (Fig. 1g, Supplementary Table 1). Recombination patterns in all driver lines appear to recapitulate the spatial, temporal and cellular gene expression patterns of the targeted endogenous loci, where such data are available (see Figures throughout the Results section). These driver and reporter lines are being deposited to the Jackson Laboratory for wide distribution. For simplicity, all drivers are named as *GeneX-CreER/Flp*, where *T2A* or *IRES* is omitted.

### Progenitor driver lines enable fate mapping of PyN developmental trajectories

The embryonic dorsal telencephalon contains a developmental ground plan whereby sequential transcription programs drive the proliferation and lineage progression of neuroepithelial and progenitor cells to orderly generate all PyNs in the cortex^3, 10, 17^ (Fig. 1c, d) Temporally regulated expression of key TF networks gates sequential developmental decisions at the progenitor, postmitotic and postnatal differentiation stages to progressively shape hierarchically organized PyN classes and subpopulations^10, 18^.

#### Probing the diversity of RGs

Among over two dozen major TFs implicated in corticogenesis^18^, the LIM-homeodomain protein LHX2 and zinc-finger transcription factor FEZF2 are foundational and act at nearly every stage throughout the construction of the neocortex (Fig. 2a). *Lhx2* plays multiple essential roles from the initial selection of cortical primordium over the hem and antihem^19–21^, to the balance of neuroepithelial proliferation over differentiation, the fate restriction of RGs, the differentiation of upper layer postmitotic PyNs^22^, and the timing of gliogenesis^17^. *Fezf2* is similarly expressed from early RGs but subsequently functions as a master regulator for the specification and differentiation of infragranular corticofugal PyNs^23, 24^. Importantly, during the period of deep layer neurogenesis, *Lhx2* suppresses *Fezf2* expression in progenitors through direct binding to its distal enhancer elements, thereby regulating the specification of PyN subtype identity^25^. However, the precise fate potential and temporal progression of *Lhx2^+^* and *Fezf2^+^* RGs (RGs*^Lhx2^*, RGs*^Fezf2^*) and their relationship to the lineage organization of cortical progenitors in general, are largely unknown. We have generated *Lhx2-CreER, Fezf2-CreER*, and *Fezf2-Flp* driver lines that enable precise fate mapping of these TF-defined RG pools.

**Figure 2:**
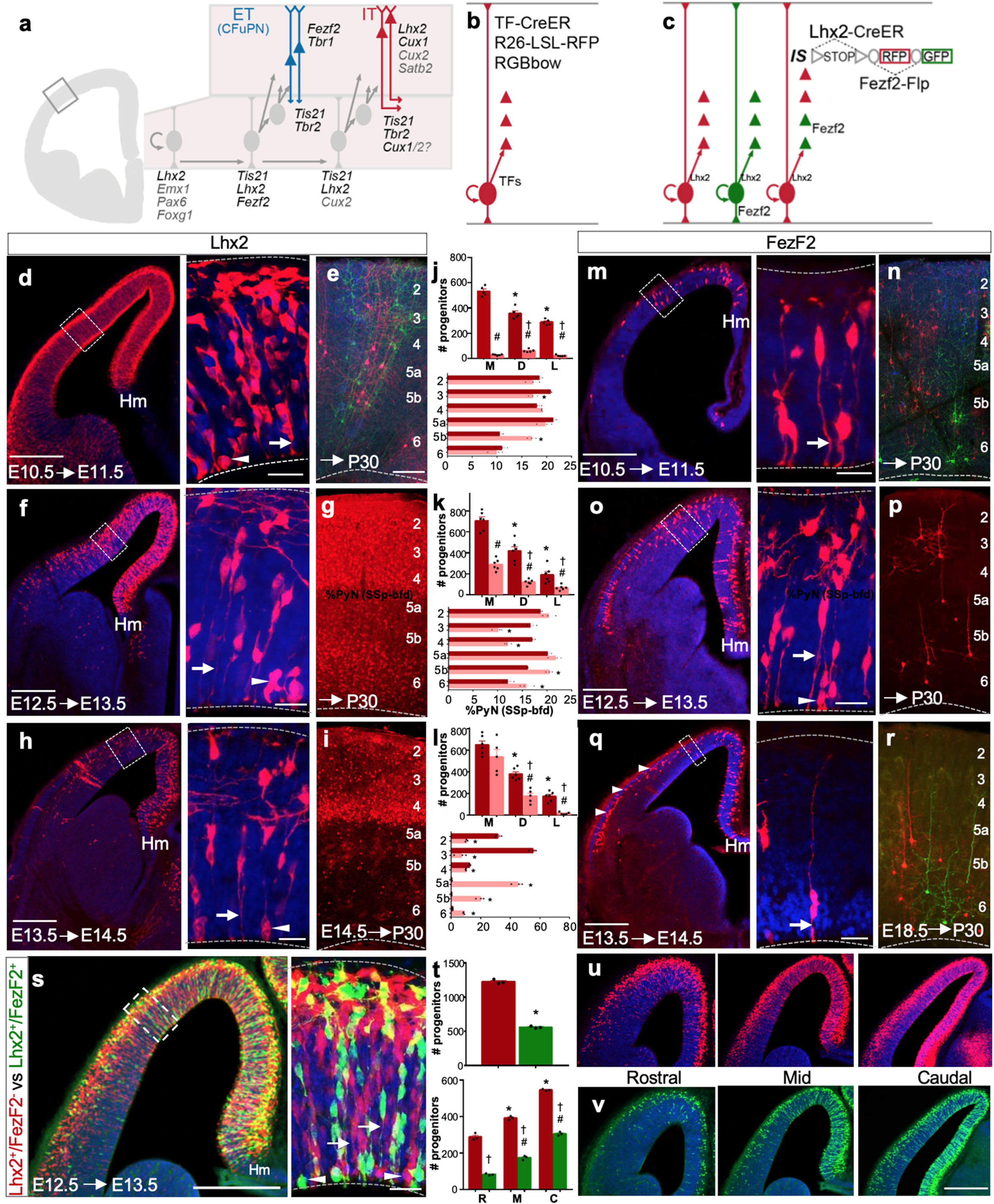
Driver lines for dissecting progenitor pools and fate mapping PyN developmental trajectories. **a**, Schematic of the lineage progression of neural progenitors of the embryonic dorsal telencephalon in a hemi-coronal section. RGs undergo either direct or indirect neurogenesis (through IPs) and give rise to PyNs of different cortical layers and projection classes. Key TFs are listed according to their expression pattern in progenitors (grey) and postmitotic neurons (red and blue); TFs used for generating driver lines are in black. **b**, Fate mapping strategy used in **d-r**: a specific TF-CreER driver labels a RG and all of its progeny when combined with a RFP or a multi-color RGBow reporter. **c**, Fate mapping scheme used in **s-v**: simultaneous labeling of different molecularly defined RGs using an intersection/subtraction reporter (IS) combined with *Lhx2-CreER* and *Fezf2-Flp* drivers; in RGs*^Lhx2+Fezf2-^*, Cre activates RFP expression only. In RGs*^Lhx2+Fezf2+^*, Cre and Flp recombinations remove the RFP cassette and activate GFP expression. **d**, 24-hour pulse-chase in a E10.5 *Lhx2-CreER;Ai14* embryo densely labeled RGs throughout the dorsal neuroepithelium; magnified view from boxed area shows RGs at different stages of the cell cycle, with endfeet (arrow) or dividing somata (arrowhead) at the ventricle wall (dashed lines). **e**, E10.5 RGs generate PyNs across layers in P30 neocortex using a strategy depicted in **b** with a *Lhx2-CreER;RGBow* mouse. **f**, Same experiment as in **d** except induced at E12.5. RGs*^Lhx2+^* distribute in a medial^high^-lateral^low^ gradient along the dorsal neuroepithelium, ending sharply at the cortex-hem boundary (Hm); magnified view shows RGs with endfeet (arrow) or dividing somata (arrowhead) at the ventricle wall. **g**, E12.5 RGs*^Lhx2+^* generated PyN progeny across all cortical layers. **h**, Same experiment as in **d** induced at E13.5. RGs*^Lhx2+^* remain distributed in a medial^high^-lateral^low^ gradient along the dorsal pallium but at reduced density compared to earlier stages. **i**, E14.5 RGs*^Lhx2+^* generated PyNs with lower density in L5/6, higher density in L2-4, and highest density in L4. **j**, Upper panel, Quantification of E10.5 RGs*^Lhx2+^* (red) (as in **d**) versus RGs*^Fezf2+^* (pink) (as in **m**) distributed across the cortical primordium, divided into medial (M), dorsal (D) and lateral (L) bins of equal length. Lower panel, laminar distribution of PyNs generated by RGs*^Lhx2+^* (red) or RGs*^Fef2+^* (pink) at E10.5, shown as percentage of total PyNs in SSp-bfd. **k**, Same quantification and comparison of RGs*^Lhx2+^* and RGs*^Fezf2+^* as in **j** except induced at E12.5. **l**, Same quantification and comparison as in **j** except RGs*^Lhx2+^* and RGs*^Fef2+^* were induced at E13.5 (upper panel). Their PyN progeny fate-mapped at E14.5 (for RGs*^Lhx2+^*) and E18.5 (for RGs*^Fezf2+^*) (lower panel). **m**, 24-hour pulse-chase in a E10.5 *Fezf2-CreER;Ai14* embryo. The spatial extent of RGs*^Fef2+^* in neuroepithelium is much restricted compared to RGs*^Lhx2+^* at this stage; magnified view shows RG endfoot (arrow) at the ventricle (dashed lines). Note the sparsity of dividing RGs. **n**, Fate mapping of E10.5 RGs*^Fef2+^* in a *Fezf2-CreER;RGBow* mouse labeled PyN progeny across all cortical layers. **o**, Same experiment as in **m** induced at E12.5. RGs*^Fef2+^* also distribute in a medial high-lateral low gradient along the dorsal neuroepithelium, ending at the cortex-hem boundary (Hm). Note that RG*^Fef2+^* density is much lower than that in **f**; magnified view shows RGs at multiple cell cycle stages with endfeet (arrow) and dividing soma (arrowhead) at the ventricle wall (dashed line). **p**, Fate mapping E12.5 RGs*^Fef2+^* using sparse labeling revealed PyNs across cortical layers. **q**, Same experiment as in **m** induced at E13.5. Only sparse RGs*^Fezf2+^* remain in medial pallium at this time, when *Fezf2* expression shifts to postmitotic PyNs (arrowheads); magnified view shows a remaining RG (arrow) at the ventricle wall (dashed line). **r**, Fate mapping at E18.5 in *Fezf2-CreER;RGBow* labeled only L5b/L6 PyNs in mature cortex. Quantification for upper panels in **j,k,l,** (n=5-6 from 2 litters): mean values, number of progenitors ± SEM. *P < 0.05 (compared with bin M, RGs*^Lhx2^*^+^), #P < 0.05 (compared with bin M, RGs*^Fezf2^*^+^), one way ANOVA, Tukey’s post hoc test. †P < 0.05 (compared with RGs*^Lhx2^*^+^ for corresponding bins), unpaired Student’s t test. Quantification for lower panels in **j,k,l**, (n=3 from 2 litters): mean values for percentage total PyNs (SSp-bfd) ± SEM. *P < 0.05 (compared with PyNs*^Lhx2^*^+^). **s**, The presence of RGs*^Lhx2+Fezf2-^* (RFP) and RGs*^Lhx2+Fezf2+^*(GFP) at E12.5 throughout the cortical primordium revealed by intersection/subtraction fate mapping schematized in **c**; magnified view shows RGs*^Lhx2+Fezf2-^* and RGs*^Lhx2+Fezf2+^* at multiple cell cycle stages with endfeet (arrow) and dividing soma (arrowhead) at the ventricle wall (dashed line). **t**, Upper panel, quantification shows that the total number of RGs*^Lhx2+Fezf2+^* (green) is approximately one-third that of RGs*^Lhx2+Fezf2-^*(red) (n=3, 2 litters). Mean values, number of progenitors ± SEM. *P < 0.0001 (compared with RGs*^Lhx2^*^+*Fezf2*-^), unpaired Student’s t test. Lower panel, quantification of RGs*^Lhx2+Fezf2-^* versus RGs*^Lhx2+Fezf2+^* (n=3, 2 litters) at rostral (R), mid-level (M) and caudal (C) sections revealed their caudal high-rostral low distribution. Mean values, number of progenitors ± SEM. *P < 0.05 (compared with rostral RGs*^Lhx2^*^+*Fezf2*-^), #P < 0.05 (compared with rostral RGs*^Lhx2^*^+*Fezf2*+^), one way ANOVA, Tukey’s post hoc test. †P < 0.05 (compared with RGs*^Lhx2^*^+/*Fezf2*-^ for corresponding regions), unpaired Student’s t test. **u** and **v**, representative sections of RGs*^Lhx2+Fezf2-^* and RGs*^Lhx2+Fezf2+^*, respectively, quantified in **t** at three rostro-caudal levels. High magnification insets are not maximum intensity projections, and do not show the z-projection seen in the low mag image. DAPI (blue) in all image panels except **e,n**. Scale bars: 20μm in **d,f,h,m,o,q,s**;100μm in all other panels.

Using the *Lhx2-CreER* driver, we carried out a series of short (∼24-hrs) pulse-chase experiments at multiple embryonic stages to reveal the status of the progenitors and lineage progression, as well as a set of long pulse-chase experiments from similar embryonic stages to the mature cortex to reveal the pattern of their PyN progeny. At E10.5, a short tamoxifen (TM) pulse-chase in *Lhx2-CreER;Ai14* embryos resulted in dense and near ubiquitous labeling of neuroepithelial cells (NEs) and RGs in the dorsal pallium at E11.5, with a sharp border at the cortex-hem boundary (Fig. 2d, j; Extended Data Fig. 1a). These early RGs showed characteristic radial fibers and end-feet at the ventricular surface; the position of their soma likely reflected the stage of cell cycle progression, including those dividing at the ventricle surface (Fig. 2d, Extended Data Fig. 1a). E12.5→E13.5 pulse-chase revealed a prominent medial^high^ to lateral^low^ gradient of RGs*^Lhx2^* (Fig. 2f,k), suggesting significant differentiation of the earlier, E10.5 RG pool. E13.5→E14.5 pulse-chase showed a similar gradient pattern at lower overall density (Fig. 2h, l, Extended Data Fig. 1c), with conspicuous cell clusters suggesting a high proliferative capacity of RGs*^Lhx2^* at this stage (Fig. 2h, Extended Data Fig. 1c). The embryonic induction patterns precisely recapitulate endogenous *Lhx2* expression patterns^26, 27^ at each stage examined (Extended Data Fig. 2a-d).

Long pulse-chase from E10.5, E12.5 and E14.5 RGs*^Lhx2^* to mature cortex labeled PyN progeny across cortical layers (Fig. 2e, g, i-l, Extended Data Fig. 1b, d), suggesting that RGs*^Lhx2^* are multipotent at these embryonic stages. E14.5 RGs*^Lhx2^*-derived PyNs show lower density in L5/6 and higher density in L2-4, suggesting more restricted fate potential (Fig. 2i, l). During the early postnatal period, when *Lhx2* expression is mostly postmitotic, pulse-chase labeled PyNs*^Lhx2^* across layers that comprise largely the IT class (Extended Data Fig. 1e). From the second postnatal week, pulse-chase labeled more astrocytes (∼60%) than PyNs, which were distributed across layers (Extended Data Fig. 1f). Together, these results indicate that the *Lhx2-CreER* driver precisely recapitulates endogenous expression across developmental stages and provides a highly valuable fate mapping tool for a TF-defined RG pool.

Similar fate mapping experiments using the *Fezf2-CreER* driver yielded a contrasting set of results. At E10.5, short pulse-chase (with the same TM dose as for *Lhx2-CreER*) labeled only a sparse set of pallial RG progenitors within the neocortex, sharply ending at the cortex-hem boundary (Fig. 2j, m, Extended Data Fig. 1g). E12.5→E13.5 pulse-chase labeled a larger set of RGs*^Fezf2^* with a similar medial^high^ to lateral^low^ gradient as RGs*^Lhx2^* but with notably lower density (Fig. 2k, o). RG*^Fezf2^* characteristics were similar to those of RGs*^Lhx2^* at the same stage (Fig. 2f, o). By E13.5, short pulse-chase labeled few RGs, mostly in the medial region, and the rest were postmitotic PyNs (Fig. 2l, q, Extended Data Figs. 1i, 2h).

Long pulse-chase from E10.5 and E12.5 to P30 cortex in *Fezf2-CreER* mice labeled PyNs distributed across cortical layers, suggesting multi-potent RGs*^Fezf2^* between E10.5-E12.5 (Fig. 2j,k,n,p; Extended Data Fig. 1h). After E13.5, Fezf2 expression is mostly shifted to postmitotic L5/6 corticofugal PyNs, especially in the lateral cortical primordium (Fig. 2l,q,r, Extended Figs. 1i, 2h). Therefore, contrasting a *Fezf2* transgenic line^14^, the *Fezf2-CreER* driver also recapitulates endogenous developmental expression (also see Extended Data Fig. 2f-h) and provides a valuable fate mapping tool for the RG*^Fezf2^* pool.

To probe the relationship between RGs*^Lhx2^* and RGs*^Fezf2^*, we first generated a *Fezf2-Flp* driver, which recapitulated endogenous *Fezf2* expression (Extended Data Fig. 2i). We then designed an intersection/subtraction (IS) strategy. Combining *Lhx2-CreER* and *Fezf2-Flp* with the *IS* reporter, we were able to differentially label RGs*^Lhx2+/Fezf2-^* (RFP) and RGs*^Lhx2+/Fezf2+^*(GFP) (Fig. 2c, s-v, Extended Data Fig. 1j). E12.5 short pulse-chase demonstrated the presence of two distinct RG subpopulations based on differential TF-expression that were intermixed across the dorsal pallium, with RGs*^Lhx2+/Fezf2-^* more than twice as abundant as RGs*^Lhx2+/Fezf2+^* (Figure 2s, t, Extended Data Fig. 1j). Both RG subpopulations were distributed in a medial^high^-lateral^low^ and rostral^high^-caudal^low^ gradient consistent with *Lhx2* and *Fezf2* expression patterns (Fig. 2f, k, o, s-v). We further analyzed the percentage of RGs*^Fezf2^* expressing LHX2 using *FezF2*-CreER embryos labeled by a 24-hour pulse-chase at E12.5. Whereas a large majority of RGs*^Fezf2^* expressed LHX2, approximately 10% did not (Extended Data Fig. 2e). Therefore, we provide compelling evidence for the presence of three distinct RG subpopulations defined by differential expression of *Lhx2* and *Fezf2*. The IS strategy can be extended to examine whether PyNs derived from RGs*^Lhx2+/Fezf2-^* and RGs*^Lhx2+/Fezf2+^* might differ in projection pattern and connectivity; this can be achieved by converting transient *CreER* activity in progenitors to a permanent toolgene expression (e.g. Flp, tTA) in progeny PyNs for viral-based phenotypic characterization and manipulation^28^.

#### Probing the diversity of neurogenic RGs (nRGs)

The early cortical progenitors comprise proliferative as well as neurogenic subpopulations, likely with different fate potentials. *Tis21* is an anti-proliferative transcription co-regulator that negatively controls cell cycle checkpoint and is specifically expressed in subsets of RGs that undergo neurogenic but not proliferative cell divisions^29, 30^. It is unknown whether the nRGs at any particular time are homogeneous or consist of subpopulations with differential TF expression and fate potential. We developed a novel fate mapping strategy to address this issue. To target nRGs, we generated a *Tis21-CreER* driver line (Extended Data Fig. 3). As *Tis21* is also expressed in neurogenic progenitors of the ventral subpallium, 48hr pulse-chase from E10.5 revealed prominent nRG-derived clones along both the pallium-glutamatergic and subpallium-GABAergic progenitor domains (Extended Data Fig. 3f). E10.5→P30 long pulse-chase revealed columnar arrangement of PyN and astrocyte clones, intermixed with subpallium-derived GABAergic interneurons (Extended Data Fig. 3g).

To restrict fate mapping to glutamatergic nRGs and probe their diversity, we designed a scheme to differentially label nRGs that either express or do not express *Fezf2*. We carried out intersection/subtraction (IS) fate mapping using *Tis21-CreER;Fezf2-Flp;IS* mice (Extended Data Fig. 3h-k). E11.5→E12.5 pulse-chase demonstrated that *Tis21/Fezf2* intersection specifically labeled a set of pallial nRG*^Fezf2^*^+^ with GFP, whereas *Tis21/Fezf2* subtraction labeled pallial and subpallial nRG*^Fezf2-^* with RFP (Extended Data Fig. 3h). The result thus revealed that pallial nRGs consisted of both *Fezf2^+^* and *Fezf2^-^* subpopulations (Extended Data Fig. 3h-k), suggestive of heterogeneity.

Long pulse-chase from E12.5→P30 revealed three types of PyN clones (Extended Data Fig. 3i-k). RFP-only clones likely derived from nRG*^Fezf2-^* in which *Tis21-CreER* activated RFP expression. They likely consisted of PyNs that did not express *Fezf2* at any stage (Extended Data Fig. 3j, j”). The GFP-only clones likely derived from nRG*^Fezf2^*^+^, in which *Tis21-CreER* and *Fezf2-Flp* co-expression activated GFP in the *IS* reporter allele (Extended Data Fig. 3k). The mixed clones containing both GFP and RFP cells are likely derived from nRG*^Fezf2-^* in which *Tis21-CreER* first activated RFP expression followed by postmitotic activation of GFP through *Fezf2-Flp* (Extended Data Fig. 3i, j, j’). Together, these results indicate the presence of nRG*^Fezf2+^* as well as nRG*^Fezf2-^*, both multi-potent in generating PyNs across all cortical layers. It remains to be examined whether their progeny might differ in terms of projection pattern and connectivity when analyzed at cellular and clonal resolution. This strategy can be used to fate map the diversity of nRGs with other TF Flp drivers.

#### Targeting intermediate progenitors

IPs and indirect neurogenesis have evolved largely in the mammalian lineage and have continued to expand substantially in primates^29, 31–33^. Along the mammalian embryonic neural tube, indirect neurogenesis is restricted to the telencephalon and is thought to contribute to the expansion of cell numbers and diversity in the forebrain, especially the neocortex. Indeed, the majority of PyNs in mouse cortex are produced through indirect neurogenesis^34–36^. But the temporal patterns of IP-mediated indirect neurogenesis in relation to their PyN progeny are not well characterized. The T-box transcription factor *Tbr2* is specifically expressed in pallial IPs throughout indirect neurogenesis^37, 38^. We have generated both *Tbr2-CreER* and *FlpER* driver lines.

Embryonic 12 hr pulse-chase at E16.5 with *Tbr2-CreER* revealed specific labeling of IPs and not RGs (Extended Data Fig. 3a, c). Long pulse-chase E16.5→P30 labeled PyNs in L2/3, which are expected to be generated at this embryonic stage (Fig. 3c, Extended Data Fig. 3d). In addition, fate mapping of *Tbr2-CreER;Ai14* from E17.5 to P28 revealed highly laminar restricted PyN cohorts in L2 (Fig. 3d). Therefore the *Tbr2-CreER* driver can achieve highly specific targeting of PyN laminar subpopulations in supragranular layers.

**Figure 3:**
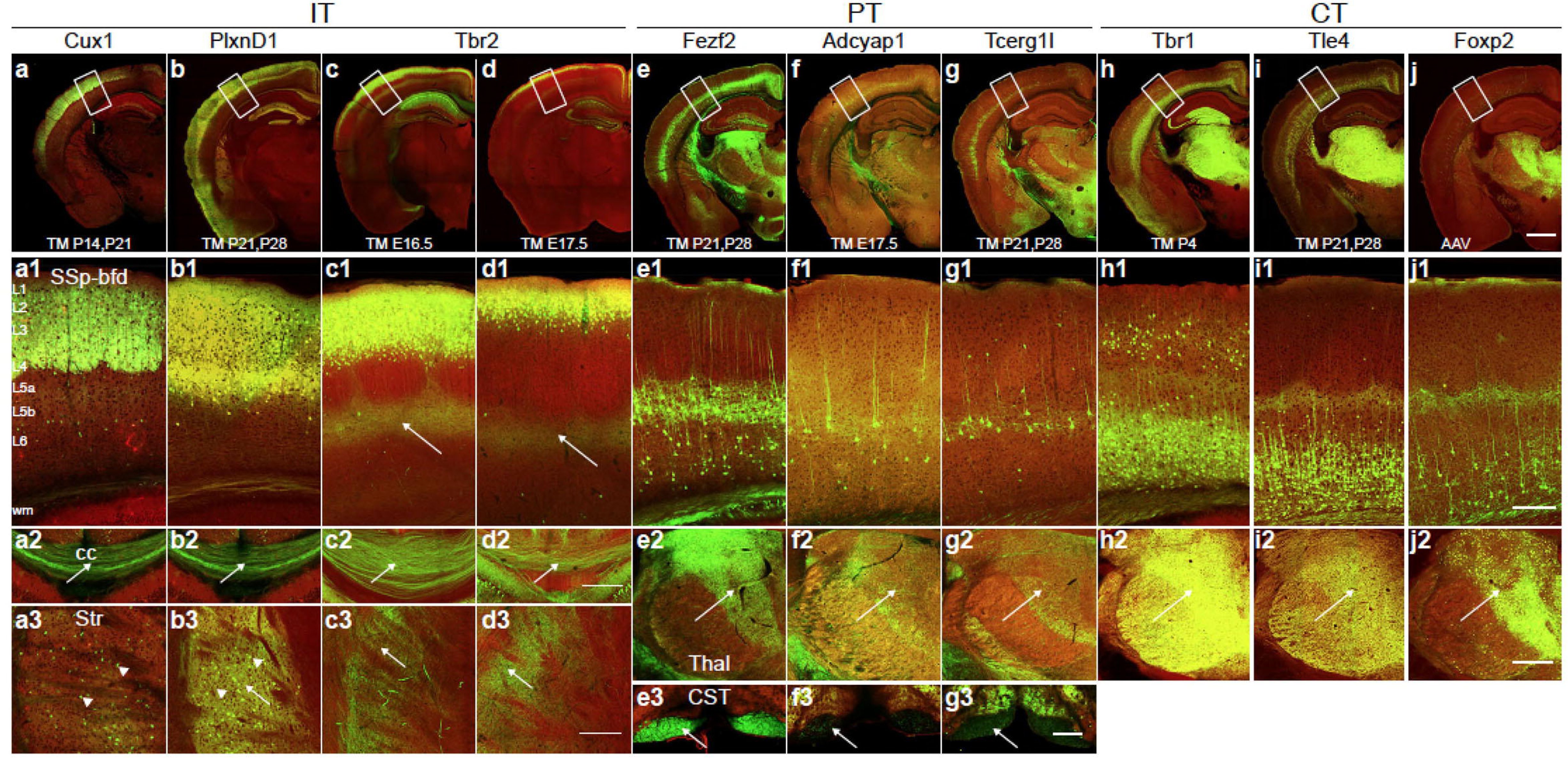
Genetic targeting of PyN projection classes and subpopulations. **a-j,** Cre recombination patterns from mouse driver lines generated and characterized in this study visualized through reporter expression (green; background autofluorescence in red). First row: coronal hemisections at Bregma −1.7 mm. **a1-d1,** IT PyN drivers targeting an array of laminar subsets in L2-5a, as visualized from a segment of the somatosensory cortex with laminar delineations, project across the corpus callosum (cc) (**a2-d2**) and to the striatum (Str) (**a3-d3**). Note *Cux1* and *PlxnD1* drivers also label subsets of medium spiny neurons in the striatum (arrowheads). *Tbr2-CreER* induction at E16 and E17 label L2/3 and L3 PyNs, respectively. Cell bodies are indicated by arrowheads and axons by arrow. **e1-g1,** PT drivers label PyNs in L5B, which project to numerous subcortical targets, including the thalamus (thal, **e2-g2**) and spinal cord (corticospinal tract–CST, **e3-g3**). cp, cerebral peduncle; POm, posterior medial nucleus of thalamus; VPM, ventral posteromedial nucleus of thalamus. **h1-j1,** CT drivers label PyNs in L6, with different thalamic nuclei as their major projection targets (**h2-j2**). TM inductions were at P21 except for *Tbr2-CreER*, as indicated, and *Adcyap1-CreER* at E17.5. The reporter allele was Ai14, except for *PlxnD1-CreER* (*Snap25-LSL-EGFP*) and *Foxp2* (systemic injection of AAV9-CAG-DIO-EGFP). Scale bars: CST panel in **g3**, 100 μm; hemisection in **j** applies to all hemisections, 1mm; all other scale bars, 200 μm.

Whereas most IPs undergo a single symmetric division to produce two PyNs, a subset are transit amplifiers, each first generating two IPs which then divide to produce a total of 4 PyNs^39^. To specifically target neurogenic IPs that also express *Tis21*, we combined *Tis21-CreER* and *Tbr2-FlpER* with the intersectional reporter Ai65. E16.5→P30 fate mapping resulted in a similar laminar pattern of PyN progeny as using *Tbr2-creER* and Ai14, consistent with the notion that most *Tbr2*^+^ IPs in the mouse dorsal pallium are neurogenic (Extended Data Fig. 3a, e). The *Tbr2-CreER* and *FlpER* driver lines thus allow efficient fate mapping and manipulation of pallial IPs.

Altogether, these progenitor driver lines enable specific and temporally controlled targeting of major pallial progenitor types (RGs, nRGs, IPs) and TF-defined subpopulations for exploring the diversity of progenitors and their lineage progression. They will facilitate tracking the developmental trajectories of PyNs from their lineage origin to postmitotic differentiation, migration, laminar deployment, circuit assembly and function.

### Driver lines targeting projection subpopulations

A useful genetic toolkit for PyNs would comprise driver lines that target their hierarchical organization^6^, including major projection classes and finer biologically meaningful subpopulations within each class. Previous efforts have produced approximately a dozen mostly transgenic lines targeting several major laminar and projection subpopulations^15, 16^. Complementary to this effort, we designed gene KI drivers based on key TFs and effector genes of the developmental genetic programs of PyNs to access more and biologically relevant subpopulations toward dissecting cortical circuit elements^10, 18, 40–42^ (Figs. 1c-g, 3). These inducible drivers further confer temporal control at different developmental stages and TM dose-dependent labeling and manipulation from sparse individual cells to dense populations. We characterized these lines with comparison to existing lines where feasible (Supplementary Tables 1, 2).

#### IT driver lines

The intratelencephalic (IT) PyNs constitute the largest top-level class and mediate the vast majority of intracortical, inter-hemispheric, and cortico-striatal communication streams^1, 43–45^. The *Cux1* and *Cux2* are Cut-repeat containing homeodomain TFs predominantly expressed in supragranular IT PyNs and their progenitors^46–48^. *Cux* expression is thought to begin at the transition from progenitor to postmitotic neurons^47, 49^, thus may initiate and then maintain the identity of IT PyNs. Although *Cux1* and *Cux2* exhibit similar DNA binding specificities and most upper layer neurons are thought to co-express these two proteins^47^, they appear to play non-redundant and additive roles in coordinating the progression of cell fate programs and in regulating dendritic branching and synapse stabilization^49, 50^. In particular, *Cux1* but not *Cux2* promotes the development of contralateral axonal connectivity of L2/3 IT cells by regulating their firing mode^51^. We generated a *Cux1-CreER* driver and compared it with the existing *Cux2* driver^12^. TM induction at P14 and P21 in *Cux1-CreER;Ai14* mice prominently labeled L2-4 PyNs largely dorsal to the rhinal fissure, as well as a set of hippocampal PyNs, recapitulating the endogenous pattern (Fig. 3a, Extended Data Fig. 4a, b, Supplementary Movie 1). Interestingly, anterograde tracing revealed that PyNs*^Cux1^* in somatosensory barrel cortex (SSp-bfd) project only to the ipsi- and contra-lateral cortex but not subcortical targets including the striatum (Fig. 4a, e, f, Supplementary Movie 2). Thus compared with current L2-4 IT drivers (e.g. Cux2^12^*, Sepw1-Cre_* NP39^15^, Rasgrf1^16^), *Cux1-CreER* represents the first cortex-restricted KI driver line and provides reliable access to a categorically distinct subpopulation of supragranular IT neurons.

**Figure 4:**
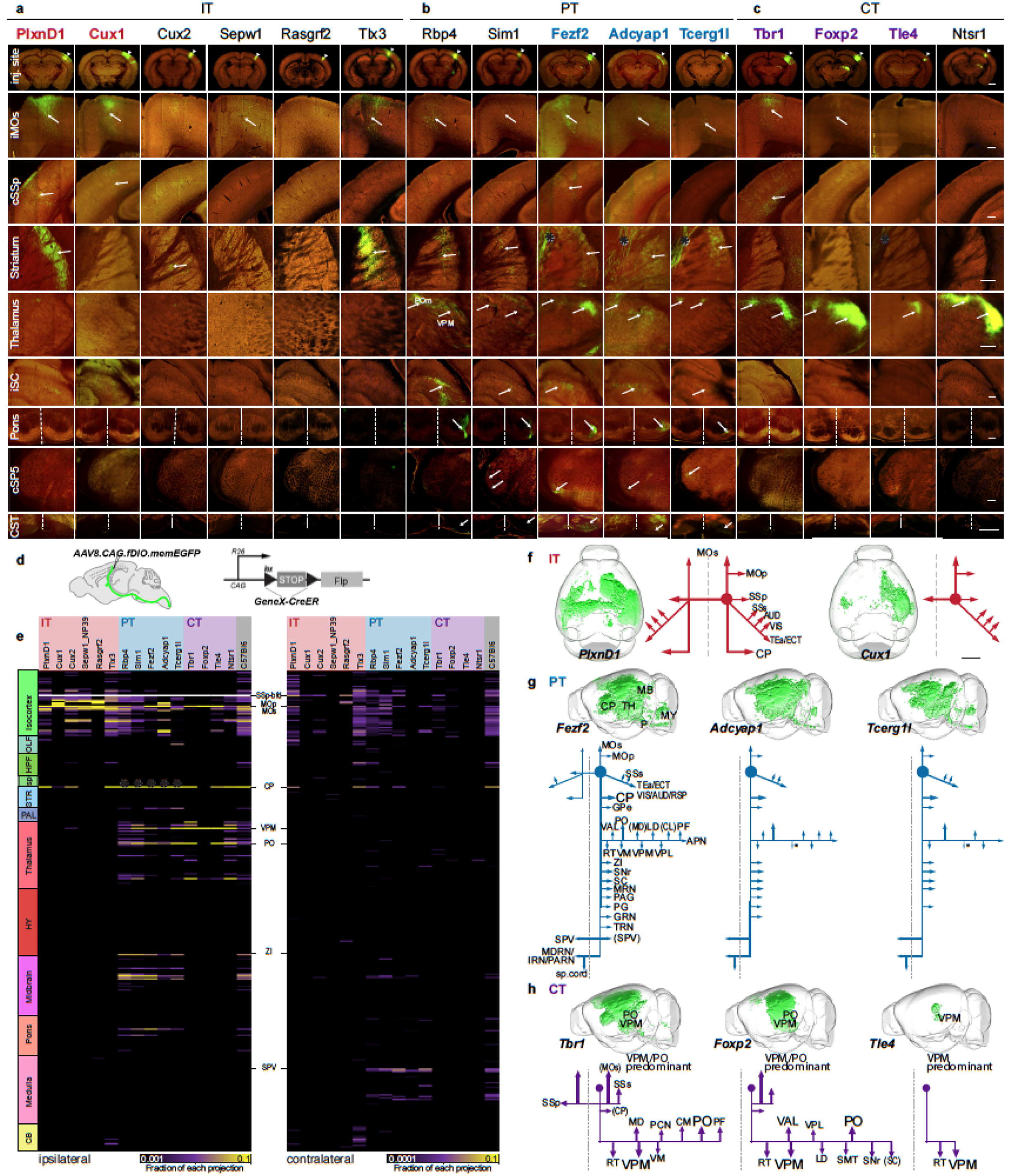
Anterograde tracing from PyN subpopulations in SSp-bfd somatosensory cortex. **a-c,** STP images at the SSp injection site (first row, arrow head) and at selected subcortical projection targets for eight driver lines characterized in this study (colored gene names) compared to seven existing driver lines (black gene names), with EGFP or EYFP expression from Cre-activated viral vector (green) and background autofluorescence (red). Arrows point to axons. **a,** IT drivers project to cortical and striatal targets. Note the absence of projection to striatum in the *Cux1* driver. **b,** PT drivers project to many corticofugal targets including brainstem and spinal cord. **c,** CT drivers project predominantly to the thalamus. **d,** Strategy for anterograde axon tracing. In a *GeneX-CreER;LSL-Flp* mouse, tamoxifen induction of CreER expression at a given time is converted to constitutive Flp expression for anterograde tracing with a Flp-dependent AAV vector. **e,** Axon projection matrix from SSp-bfd to 321 ipsilateral and 321 contralateral targets (in rows), each grouped under 12 major categories (left boxes), for each of the mouse lines presented in **a-c**. Color shades in each row represent the fraction of total axon signals measured from a single experiment per brain area. Signals in the injection site was subtracted from the whole brain to show fraction of projections outside the injection site (appears in white). PyNs*^PlxnD1^* project bilaterally to cortex and striatum; PyNs*^Cux1^* project bilaterally to cortex but not to striatum; PyNs*^Fezf2^*, PyNs*^Adcyap1^* and PyNs*^Tcerg1l^* project to multiple ipsilateral targets and to the contralateral brainstem (arrows); and PyNs*^Tbr1^* project bilaterally to cortex and to ipsilateral thalamus, PyNs*^Foxp2^* and PyNs*^Tle4^* project to the ipsilateral cortex and thalamus. **f-h,** Schematics of main projection targets for each PyN subpopulation generated in this study and whole-brain three dimensional renderings of axon projections registered to the CCFv3 for each PyN subpopulation in the SSp-bfd. Scale bars: first row in **c** (applies to first row), 1 mm; second to eighth rows in **c** (applies to each respective row), 200 μm; CST panel (bottom row) in **c** applies to entire row, 100 μm; **h** (applies to all 3D renderings in f-h), 2 mm. Asterisks in **b, c, e** and **g** indicate presence of passing fibers.

The supragranular layers comprise diverse IT types that project to a large combination of ipsi- and contra-lateral cortical and striatal regions^42, 44, 52^, but only a few L2/3 driver lines have been reported to date^16^, and none distinguish L2 vs L3 PyNs. We used a fate mapping approach to target L2/3 PyNs by engaging cell lineage and birth timing. In our *Tbr2-CreER* driver targeting IPs (Fig. 2q), TM induction at E16.5 and E17.5 specifically labeled PyNs L2/3 and L2, respectively (Fig. 3c, d). Combined with the CreER→Flp conversion strategy that converts transient lineage and birth timing signals to permanent Flp expression^28^, this approach enables specific AAV manipulation of L2 and L3 IT neurons.

The *plexinD1-semaphorin3E* receptor-ligand system has been implicated in neuronal migration, axon guidance, synapse specification^53, 54^, as well as vascular development^55^. In developing and mature cortex, *PlxnD1* is expressed in large sets of callosal projection IT PyNs (CPNs) at especially high levels during the early postnatal period of synapse maturation and refinement^40, 52^. We generated *PlxnD1-CreER* and *PlxnD1-Flp* lines. Both drivers recapitulated endogenous *PlxnD1* expression and labeled projection neurons in the cerebral cortex, hippocampus, amygdala (BLA and LA), and medium spiny neurons of the striatum (Figs. 3b, 6e, Extended Data Figs. 4, 5a-c, Supplementary Tables 3, 4, Supplementary Movies 3, 4). In the neocortex, L5A and L2/3 IT PyNs were labeled (Fig. 3b, Extended Data Fig. 4a, d), with their axons predominantly restricted to the cortex and striatum (Fig. 4a). In the piriform cortex, L2 PyNs were targeted in both *PlxnD1-CreER* and *PlxnD1-Flp* lines (Supplementary Movies 3, 4). As *PlxnD1* is also expressed in a set of vascular cells, we bred *PlxnD1-CreER* with the neuron-specific reporter *Snap25-LSL-EGFP* to achieve selective labeling of *PlxnD1^+^* PyNs (PyNs*^PlxnD1^*, Fig. 3b).

**Figure 5:**
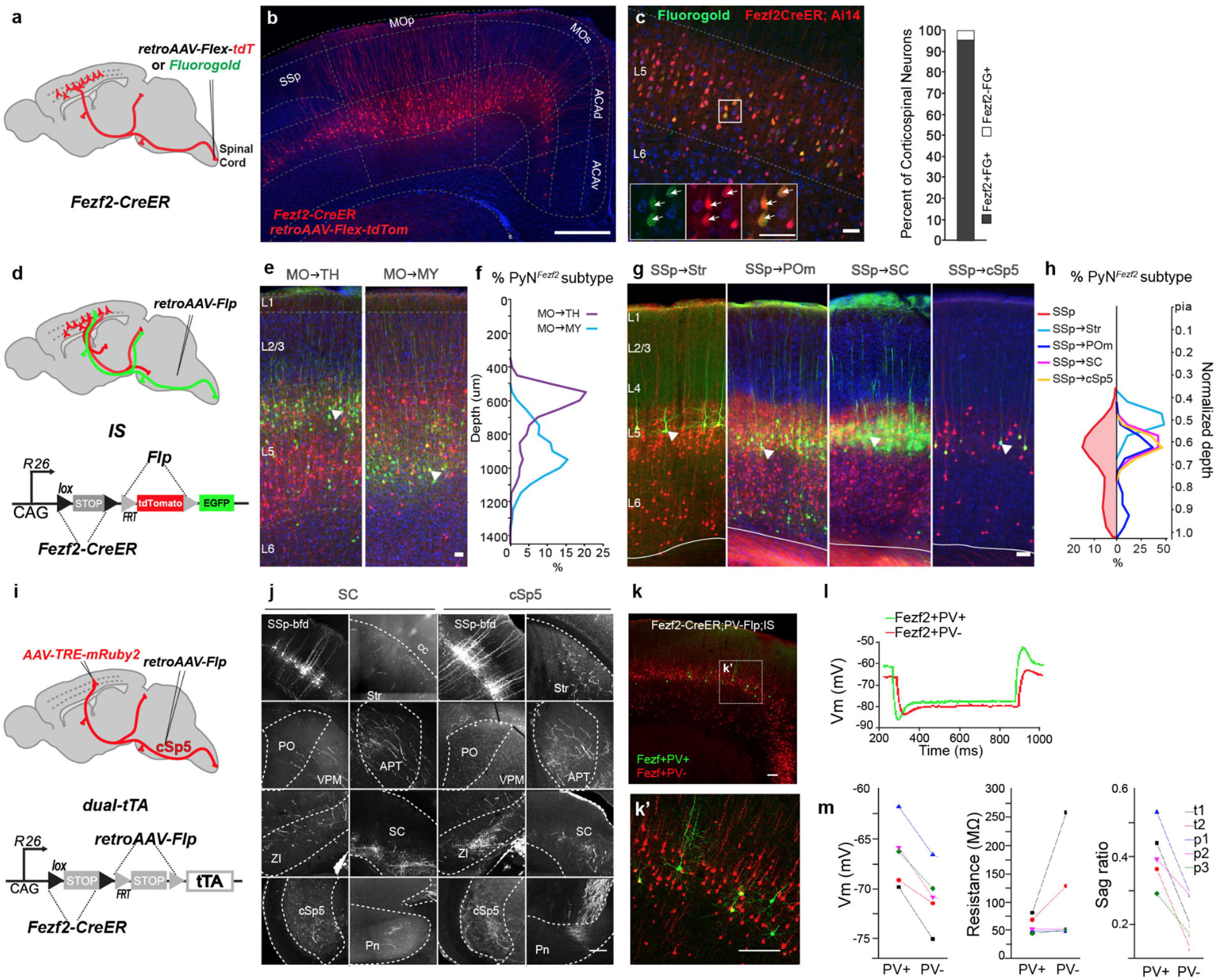
Intersectional dissection of PyN*^Fezf2^* subpopulations. **a,** Strategy for retrograde labeling of PyN*^Fezf2^* subpopulations by viral and tracer injections from the spinal cord of *Fezf2-CreER* mice. **b,** PyN*^Fezf2^* corticospinal neurons in L5B of sensorimotor cortex labeled by retroAAV-flex-tdT. **c,** *Fezf2CreER;Ai14* captures >95% of fluorogold retrograde labeled corticospinal neurons. Inset arrows show *Fezf2* and fluorogold co-labeling. **d,** An intersection-subtraction (IS) reporter strategy to label projection-defined PyN*^Fezf2^* subpopulations (retroAAV-Flp, GFP) among the whole population (*Fezf2-CreER*, RFP). **e,f,** In the motor cortex (MO), thalamus-projecting PyNs*^Fezf2^* are located in the upper L5B, whereas the medulla-projecting PyN*^Fezf2^* subpopulation are located in lower L5B. **g,** PyNs*^Fezf2^* labeled from a defined projection target (GFP) show more restricted sublaminar position in L5 compared to the whole population (RFP) in SSp cortex. **h,** Normalized cortical depth distributions of overall PyN*^Fezf2^* population (leftward curve) and of each target-defined subpopulation (rightward curves) in SSp. **i,** Strategy for combinatorial targeting by marker, axon target and soma location (‘triple trigger’). The dual-tTA reporter expresses the transcription activator tTA upon Cre and Flp recombination. In a *Fezf2-CreER;dual-tTA* mouse, TM induction combined with retroAAV-Flp injection at the contralateral spinal trigeminal nucleus (cSp5) activate tTA expression in cSp5-projection PyNs^Fezf2^ across cortical areas, and AAV-TRE-mRuby injection at SSp-bfd then labels cSp5-projection PyNs^Fezf2^ in the SSp-bfd. **j,** Coronal images of PyNs^Fezf2^ in SSp-bfd with projection to SC or cSp5, displaying axon collaterals at various subcortical targets, including Str, ZI, thalamus, pons. **k,** Using the *Fezf2-CreER;Pv-Flp;IS* triple allele mice, PV^-^ and PV^+^ PyNs^Fezf2^ can be distinguished by their expression of RFP and GFP, respectively. Boxed area is magnified in k’. **l,** Sample voltage responses induced by current injection from a pair of PV^+^ (green) and PV^-^ (red) PyNs^Fezf2^ by whole-cell patch recording in a cortical slice. **m,** Electrophysiological differences between 5 pairs of PV^+^ and PV^-^ PyNs^Fezf2^: resting membrane potential (Vm, −66.5 ± 1.6 vs. −70.7 ± 1.5 mV, mean ± s.e.m.; p = 0.0014, Student’s paired t-test); input resistance (MΩ, 60.1 ± 7.8 vs. 108.7 ± 45.4 MΩ, mean ± s.e.m.; p = 0.23, Student’s paired t-test); Sag ratio (=hyperpolarization (Peak-Steady)/Steady, 0.40 ± 0.05 vs. 0.21 ± 0.04, mean ± s.e.m.; p = 0.0039, Student’s paired t-test). Abbreviations: cc, corpus callosum; SSp-bfd, somatosensory barrel field cortex; MO, motor cortex; APT, anterior pretectal nucleus; Str, striatum; SC, Superior Colliculus; PO, posterior nucleus of the thalamus; VPM, ventral posteromedial nucleus of the thalamus; cSp5, contralateral spinal trigeminal nucleus; PV, Parvalbumin. Scale bars: b, 500μm; c,e,g, 50μm; j, 200μm; k, 100μm.

**Figure 6:**
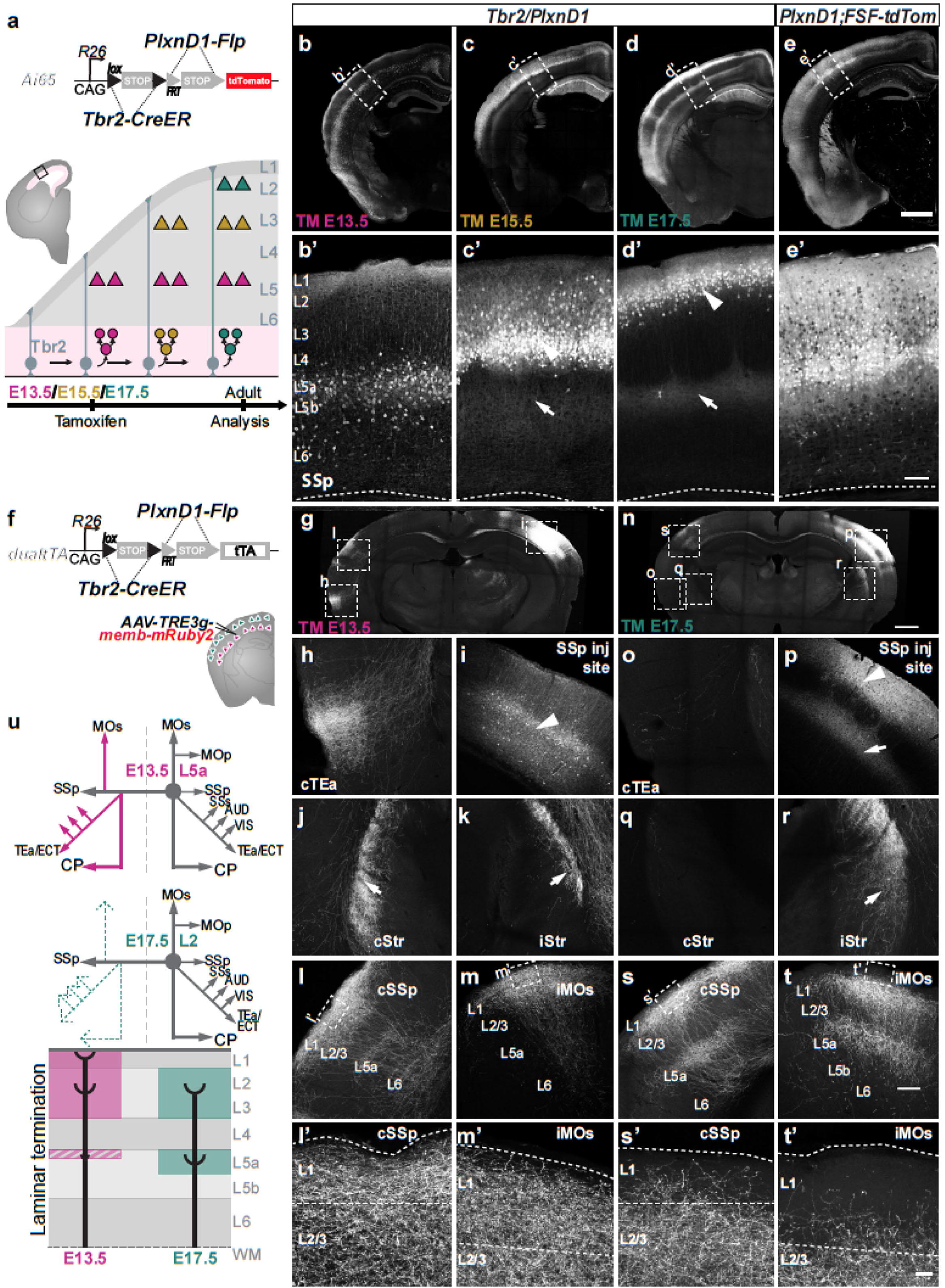
Combinatorial targeting of PyN subtypes defined by lineage, birth time, anatomical location, projection pattern. **a,** Strategy for combinatorial labeling of PyNs*^PlxnD1^* laminar subsets. In a *Tbr2-CreER;PlxnD1-Flp;Ai65* mouse, TM inductions at successive embryonic times label PyNs*^PlxnD1^* born from intermediate progenitors at these times that occupy deeper and more superficial layers. **b-d,** Representative images of laminar subsets born at E13.5 (b), E15.5 (c), and E17.5 (d). **e,** As a comparison, the whole PyNs*^PlxnD1^* population is labeled in a *PlxnD1-Flp;R26-CAG-FSF-tdTom* mouse. **b’-e’**, High magnification of boxed regions in b-d. Arrowheads indicate cell body positions; arrows indicate axons. **f,** In a *Tbr2-CreER;PlxnD1-Flp;dual-tTA* mouse, tamoxifen inductions at E13.5 and E17.5 activates tTA expression in L5A and L2 PyNs*^PlxnD1^*, respectively; AAV-TRE3g-memb-mRuby2 injection in barrel cortex then reveals the axon projection pattern of each laminar specific PyNs*^PlxnD1^*. **g-m**, Anterograde tracing from E13.5-born L5A PyNs*^PlxnD1^* in SSp-bfd. **g,** Coronal section shows AAV-TRE3g-memb-mRuby2 injection site and several major projection targets. **h-m,** Example images of axon projection targets of several indicated ipsi- and contralateral sites. Arrowheads indicate somata; arrows indicate axons. (**n-t**) Anterograde tracing from E17.5-born L2 PyNs*^PlxnD1^* in SSp-bfd. **n,** Coronal section shows AAV injection site and several major projection targets. **o-s,** Example images of axon projection targets of several indicated ipsi- and contra-lateral sites. **l’,m’,s’,t’,** Higher magnification of cSSp (**l’ & s’**) and iMOs (**m’ & t’**) display laminar axon termination differences between L5A versus L2 PyNs*^PlxnD1^*, respectively. **u**, Schematic comparison of the projection patterns of E13.5-born L5A versus E17.5-born L2 PyNs*^PlxnD1^*. Note that L5A and L2 PyNs*^PlxnD1^* show major differences in the strength of several contralateral targets and in the laminar pattern of axon termination. Scale bar in: **e** (applies to **b-e**) & **n** (applies to **g & n**), 1 mm; **e’** (applies to **b’-e’**), 200 μm; **t** (applies to **h-t**), 50μm; **t’** (applies to **l’, m’, s’ & t’**), 5μm.

Cell type specific anterograde tracing using a CreER→ Flp conversion strategy (Fig. 4d) and Flp-activated AAV-fDIO-EGFP from whisker SSp-bfd reveals that PyNs*^PlxnD1^* project to ipsi- and contra-lateral cortical and striatal regions (Fig. 4a, e, f, Extended Data Fig. 7a, b, o, s, t, Supplementary Tables 5-7, Supplementary Movie 5). Thus, the *PlxnD1-CreER* and *PlxnD1-Flp* drivers provide versatile experimental access to this major IT subpopulation and to the *PlxnD1^+^* subpopulations in the striatum and amygdala. Together, the new IT driver lines reported here are more specific, robust, and reliable in targeting IT neurons compared with existing tools (Supplementary Table 2).

#### PT drivers

Following its early expression in a subset of dorsal pallial progenitors, *Fezf2* becomes restricted to postmitotic L5 and L6 corticofugal PyNs, with high levels in L5B PT and low levels in a subset of CT neurons^40, 56^. At postnatal stages, *Fezf2-CreER* and *Fezf2-Flp* drivers precisely recapitulate endogenous postmitotic *Fezf2* expression (Fig. 3e, Extended Data Figs. 4, 5d-f, Supplementary Movie 6). *Fezf2-CreER;Ai14* labeled projection neurons in the cerebral cortex, hippocampus, amygdala, olfactory bulb and sparsely in the cerebral nuclei (striatum, nucleus accumbens and olfactory tubercle) (Extended Data Fig. 4c, Supplementary Table 4, Supplementary Movie 6). Within the neocortex, PyNs*^Fezf2^* are predominantly located in L5B and to a lesser extent in L6 (Fig. 3e1, Extended Data Figs. 4a, d, 5d-f). There was a conspicuous lack of expression in the evolutionarily more ancient cortical regions below the rhinal fissure (Supplementary Table 4, Supplementary Movie 6). Anterograde tracing of PyNs*^Fezf2^* in SSp-bfd revealed projections to numerous somatomotor cortical (e.g. ipsilateral vibrissal MOs) and subcortical regions including striatum, VPM and POm in the thalamus, anterior pretectal nucleus, ipsilateral superior colliculus (iSC), pontine nucleus, cortiscospinal tract (CST) and contralateral spinal trigeminal nucleus (cSP5) (Fig. 4b, e, g, Extended Data Fig. 7a, b, h-n, Supplementary Tables 5, 6, Supplementary Movie 7).

In addition to the *Fezf2* drivers targeting the broad corticofugal class, we further generated several lines targeting subpopulations. *Adcyap1* encodes a neuropeptide and is a transcriptional target of *FEZF2*^57^, expressed in a subset of PyNs*^Fezf2^* located in upper L5B mainly during the postmitotic and perinatal stage. Late embryonic TM induction in *Adcyap1-CreER* recapitulated endogenous expression pattern (Fig. 3f, Extended Data Figs. 4, 5d-f, Supplementary Table 4, Supplementary Movie 8). Anterograde tracing of PyNs*^Adcyap1^* revealed more restricted projections to ipsilateral vibrissal motor cortex (MOs), striatum, thalamus (predominantly POm), iSC, pons, CST and cSP5, overlapping with a subset of PyN*^Fezf2^* targets (Fig. 4b, e, g, Extended Data Fig. 7a, b, Supplementary Tables 6, 7, Supplementary Movie 9).

*Tcerg1l* is a transcription elongation regulator expressed in a subset of L5 corticofugal PyNs during postmitotic and early postnatal stages^40^. Single cell analysis of molecular genetic trajectories across cortical neurogenesis revealed that *Tcerg1l* defines a distinct sub-branch of PyNs*^Fezf2^* postmitotic differentiation^58^. TM induction at P21 in *Tcerg1l-CreER* labeled a subset of L5B PyNs, and also neurons in the septum, hypothalamus, midbrain (PAG) and retina (Fig. 3g, Extended Data Figs. 4, 5d-f, Supplementary Movie 10). Anterograde labeling of PyNs*^Tcerg1l^* revealed more restricted projections compared to those of PyNs^Fezf2^, such as an absence of collaterals to first-order thalamic nucleus VPM (Fig. 4b, e, g, Extended Data Fig. 7a, b, Supplementary Tables 5, 6, Supplementary Movie 11). We expect that future studies (e.g. single cell reconstruction) using this line may reveal more specific subtype(s) of PT cells.

SEMA3E engages in ligand/receptor interactions with PLXND1 to regulate axon guidance and synapse specificity^55^. *Sema3E* is expressed in a subset of L5B PyNs primarily in the sensory areas ^53, 59^. The *Sema3E-CreER* driver ^60^ labels a subset of L5B PyNs (Extended Data Fig. 4d; Supplementary Table 4, Supplementary Movie 12). Anterograde labeling revealed that PyNs*^Sema3E^* projected to more restricted subcortical areas, namely higher order thalamic nucleus POm and pontine nucleus (Extended Data Fig. 7c-g).

Among the two more frequently used L5 transgenic Cre lines, *Rbp4_Kl100* includes an unknown mixture of PT and IT cells^15^, and *Sim1_KJ18* is not well expressed in posterior cortex (Extended Data Fig. 4a). The new set of KI driver lines reported here cover the vast majority of PT neurons across neocortical areas and more restricted subsets, thus together they enable a finer hierarchical dissection of molecularly and anatomically defined PT types.

#### CT driver lines

The T-box transcription factor *Tbr1* is thought to represent an obligatory step of the TF cascade of PyN neurogenesis and to express in most if not all newly postmitotic PyNs^35^. *Tbr1* then becomes mainly restricted to L6 CT neurons and directly represses the expression of *Fezf2* and *Ctip2* to suppress the SCPN (PT) fate^10, 61^. Embryonic TM induction at E14.5 and E15.5 revealed that Tbr1 remained restricted to a subset of L6 CT and a small proportion of L2-4 PyNs, and was not expressed in all newly postmitotic PyNs at these stages, which would have been mostly deployed to L2/3 (Extended Data Fig. 6). Therefore, our result contrasts the previous notion based on immunolabeling that *Tbr1* is an obligatory step in all newly born PyNs following a sequence of TF expression of *Pax6*, *Tbr2* and *Tbr1* in RGs, IPs and postmitotic PyNs, respectively^37^. TM induction at P4 in the *Tbr1-CreER* driver labeled dense L6 CT neurons, with sparse labeling in L2/3, as well as subsets of cells in OB, cerebral nuclei (i.e. pallidum), hippocampus (i.e. dentate gyrus), piriform cortex, and sparse expression in amygdala (Figure 3h; Extended Data Fig. 4a-c, Supplementary Table 4; Supplementary Movie 13). Anterograde tracing revealed that PyNs*^Tbr1^* projected to multiple thalamic targets, including primary and higher order nuclei (e.g. VPM, VM, POm, MD), as well as RT (Fig. 4c, e, h, Extended Data Fig. 7a, b, Supplementary Tables 5, 6, Supplementary Movie 14). Consistent with a more recent study showing that a subset of L6 *Tbr1* PyNs also project to the contralateral cortex^62^, we found that the corpus callosum was labeled in adult *Tbr1-CreER;A14* mice induced at E14.5, E15.5 and P4 (Ext data Fig. 6). It remains to be determined whether the set of L6 and/or L2/3 PyNs*^Tbr1^* that contributed to the contralateral cortex projection (Fig. 4e, h) might represent distinct PyN types.

*Transducin-Like Enhancer Of Split 4* (*Tle4*) is a transcription corepressor expressed in a subset of CT PyNs^40, 63^. Our *Tle4-CreER* driver specifically labels L6 CT PyNs across the cortex (Fig. 3i, Extended Data Figs. 4, 5d-f; Supplementary Table 4, Supplementary Movie 15). *Tle4* is also expressed in medium spiny neurons of the striatum, OB, hypothalamus, iSC, cerebellum, septum and sparsely in the hippocampus, amygdala and brainstem (Extended Data Fig. 4c, Supplementary Table 4, Supplementary Movie 15). Anterograde tracing revealed that PyNs*^Tle4^* in SSp-bfd projected specifically to first order thalamic nucleus VPM and RT (Fig. 4c, e, h, Extended Data Fig. 7a, b, Supplementary Table 4, Supplementary Movie 16).

The *forkhead/winged helix transcription factor, Foxp2*, is expressed in many CT neurons from postmitotic stage to mature cortex, and transiently expressed in a minor subset of ET neurons in the first postnatal week^64, 65^. We characterized a *Foxp2-IRES-Cre* KI line^66^ by systemic injection of Cre dependent AAV9-DIO-GFP in 2 month old mice to reveal its brain-wide cell distribution pattern. Cortical labeling was restricted to L6 PyNs, and *Foxp2*^+^ cells were found in multiple other brain regions, including striatum, thalamus, hypothalamus, midbrain (periaqueductal gray, superior and inferior colliculi), cerebellum and inferior olive (Fig. 3j, Extended Data Fig. 4a-c, Supplementary Table 4, Supplementary Movie 17). Anterograde labeling revealed that PyN*^Foxp2^* in SSp-bfd projected to thalamus, tectum and some ipsilateral cortical areas (Fig. 4c; Supplementary Table 4; Supplementary Movie 18). Compared to PyNs*^Tle4^*, PyN*^Foxp2^* projected more broadly to the thalamus, largely overlapping with the thalamic targets of PyNs*^Tbr1^*.

In comparison with the *Ntsr1* transgenic line, ∼90% of L6 Tle4*^+^* or *Ntsr1-Cre* labeled cells are *Foxp2^+^* in the somatosensory cortex^65, 67^. The new CT driver lines have broader coverage of cortical areas (Ext data Fig. 4b), and broader (*Tbr1, Foxp2* drivers) as well as more specific (*Tle4* driver) coverage of the CT projection types (Fig. 4h).

To further validate and characterize several PyN driver lines, we carried out a set of histochemical analysis using a panel of class-defining markers, specifically CUX1, BRN2, or SATB2 (restricted to IT); CTIP2, LDB2 (enriched in PT); FOG2 (restricted to CT). *Tbr2-*, *Lhx2-*, *Cux1-*, and *PlexinD1*-CreER labeled different laminar subsets of PyNs that almost exclusively co-expressed IT specific markers. PyNs derived from *FezF2*-CreER extensively co-labeled with PT specific markers, and those derived from *Tcerg1l-* and *Adcyap1*-CreER also co-labeled with PT markers. PyNs generated from *Tle4*- and *Tbr1*-CreER co-labeled a set of CT markers. Interestingly, *FezF2*-CreER derived PyNs also labeled some L6 CTIP2^+^ CT PyNs, while *Tle4*-CreER derived PyNs labeled some L5b cells expressing PT-specific markers (Ext Data Fig. 5). The laminar patterns and class-specific marker expression in all these driver lines precisely recapitulated endogenous patterns (Ext data Fig 8, <u>Reference Atlas :: Allen Brain Atlas: Mouse Brain (brain-map.org)</u>), further demonstrating the reliability and specificity of these driver lines.

In summary, compared with existing tools^15, 16^, the new set of KI lines target the hierarchical organization of PyNs, are more specific, reliable and stable, and further allow inducible and combinatorial targeting (Supplementary Tables 1, 2, 4). Furthermore, by combining a CreER and a Flp allele into the same mouse, we were able to differentially target PyNs*^Fezf2^* and PyNs*^Tle4^* (Extended Data Fig. 7h-n) or PyNs*^Fezf2^* and PyNs*^PlxnD1^* (Extended Data Fig. 7o-t) in the same mouse. Importantly, most KI lines cover extensive cortical and neocortical areas consistent with endogenous gene expression (Extended Data Fig. 4), suggesting that the areal patterns of cells expressing the marker gene are faithfully covered by the driver line; this feature enables investigators to study many specific areas across global cortical networks with cell type resolution. In contrast, most transgenic lines show partial expression across cortical areas (e.g. *Sim1, Ntsr1*; Extended Data Fig. 4, Supplementary Table 4), which preclude their use in non-expressing areas.

### Combinatorial genetic and retrograde viral targeting of projection types

Our anterograde tracing results demonstrate that most marker-defined PyN subpopulations project to multiple brain regions (Fig. 4; Extended Data Fig. 7). To examine whether each subpopulation may comprise multiple projection types with more restricted targets and laminar location, we used retrograde AAV (retroAAV) and combinatorial approaches to trace from a defined target in a few driver lines (Fig. 5, Extended Data Fig. 8).

The *Fezf2-CreER* driver efficiently captured the broad PT population (Fig. 5a-c, Extended Data Fig. 8, Supplementary Table 6). retroAAV-Flex-tdTomato injection in the spinal cord of *Fezf2-CreER* mice specifically and prominently labeled L5B corticospinal PyNs in the sensory and motor cortex, e.g. caudal sensorimotor forelimb areas (Fig. 5b), MOs, rostral forelimb area and SSs (not shown). Fluorogold injection in the cervical spinal cord of *Fezf2-CreER;Ai14* mice revealed that over 95% of spinal-projecting PyNs in MO were *Fezf2^+^* (Fig. 5c). To explore whether the broad PyNs*^Fezf2^* in MO or SSp-bfd comprise subpopulations jointly defined by projection targets and sub-laminar position, we used the IS reporter^28^. In *Fezf2-CreER;IS* mice, *Cre* recombination labeled the broad PyNs*^Fezf2^* with RFP, while injection of retroAAV-Flp activated GFP expression in the subset of PyNs*^Fezf2^* projecting to the injection target (Fig. 5d). Consistent with a previous finding^68^, retroAAV tracing from the thalamus and medulla labeled PyNs*^Fezf2^* in the upper and lower sublamina of L5B, respectively (Fig. 5e, f, Extended Data Fig. 8c), suggesting distinct PyNs*^Fezf2^* subtypes in the MO. In the SSp-bfd, retrograde labeling revealed that PyNs*^Fezf2^* with collaterals to the striatum were located in upper L5, those with collaterals to superior colliculus or contralateral spinal trigeminal nucleus (Sp5) in the brainstem were restricted to middle and lower portion of L5B, and those projecting to the POm thalamus were located in both the middle to lower portion of L5B as well as L6 (Fig. 5g, h, Extended Data Fig. 8e-h).

We further used retrograde labeling to dissect the CT and IT populations. In the *Tle4-CreER;IS* mice, retrograde tracing from the VPM thalamus revealed two subpopulations of L6 PyNs*^Tle4^*, one extended apical dendrites to L4/5 border while the other to L1 (Extended Data Fig 8i), suggesting two subpopulations with differential inputs. In the *PlxnD1-CreER;IS* mice (Extended Data Fig 8d) and *PlxnD1-CreER* mice (Extended Data Fig 8j-o), retrograde labeling from the striatum revealed that while L5A PyNs*^PlxnD1^* projected to both ipsi- and contra-lateral striatum, L2/3 PyNs*^PlxnD1^* projected mostly to the ipsilateral striatum.

In addition, we designed a novel strategy (triple trigger) to target PyNs jointly defined by a marker (driver line), a projection target, and a cortical location. We generated a new intersectional reporter, *dual-tTA*, that converts Cre and Flp expression to tTA expression for AAV manipulation (Fig. 5i). In a compound *Fezf2-CreER;dual-tTA* mouse, retroAAV2*-Flp* injection in the superior colliculus or contralateral Sp5 activated tTA expression in SC- or Sp5-projecting PyNs*^Fezf2^* in the cortex, and AAV-TRE-mRuby to SSp-bfd then labeled these restricted projection types (Figure 5j). As a control, we injected *AAV-TRE-mRuby* to SSp-bfd without retroAAV-Flp, ensuring that *dual-tTA* expression was dependent on both Cre and Flp (Extended Data Fig. 8q).

Using an orthogonal approach to further reveal the diversity of L5B PyNs*^Fezf2^*, we compared subsets that either express or do not express the calcium binding protein parvalbumin. Using the *Fezf2-CreER;Pv-Flp;IS* compound allele mice, we differentially labeled PyNs*^Fezf2+/PV-^* (RFP) and PyNs*^Fezf2+/PV+^* (EGFP) and compared their intrinsic properties using patch clamp recordings in cortical brain slices (Fig. 5k-m). We found that, compared to PyNs*^Fezf2+/PV-^*, PyNs*^Fezf2+/PV+^* exhibited more depolarized resting membrane potentials and a larger hyperpolarization-activated depolarization. This result suggests that PV^+^ PyNs*^Fezf2^* are more excitable and play a stronger rebound activity after hyperpolarization (Fig. 5m). Whether these subpopulations have distinct subcortical projection targets remains to be determined.

Finally, our anterograde labeling of PyNs*^Fezf2^* revealed a set of unexpected contralateral cortical and striatal projections (Fig. 4e, g). To explore this PyNs*^Fezf2^* subpopulation, we noted that our retrograde CTB tracing from the striatum also labeled a set of contralateral PyNs*^Fezf2+^* located at the L5A/L5B border of SSp-bfd and MO (Extended Data Fig. 9a-d), a feature characteristic of IT cells. To explore the molecular identity of this subpopulation, we performed a set of double mRNA in situ hybridization experiments and found that a small set of PyNs located at the L5A/L5B border region co-expressed *Fezf2* and *PlxnD1* (Extended Data Fig. 9e). Using an *IS* strategy, we confirmed that this PyNs*^Fezf2/PlxnD1^* subpopulation occupied the very top sublayer of the broad PyN*^Fezf2^* population (Extended Data Fig. 9f). Therefore, PyNs*^Fezf2/PlxnD1^* likely contributed to the contralateral cortical and striatal projection of the PyNs*^Fezf2^* population (Fig. 4e, g). These results suggest that the role *Fezf2* in postmitotic differentiation of ET neurons might include the generation of a small subpopulation with IT features. Single cell reconstruction may reveal whether PyNs*^Fezf2/PlxnD1^* are typical IT cells or also project subcortically as most other PyNs*^Fezf2^* and represent an “intermediate PT-IT hybrid type”.

### Combinatorial targeting of PyN subtypes by lineage, cell birth order, marker and anatomy

Compared with the PT and CT classes which are restricted to infragranular layers and generated during the early phase of cortical neurogenesis, IT is a particularly diverse class comprising PyNs across layers that are generated throughout cortical neurogenesis, likely with numerous combinatorial projection patterns to many dozens of cortical and striatal targets. For example, the PyN*^PlxnD1^* population as a whole localizes to L5A, L3 and L2 and projects to many ipsilateral and contralateral cortical and striatal targets (Figs. 3b, 4a, e, f, 6, Extended Data Figs. 4, 5, Supplementary Tables 1, 4-7). It is unknown whether PyNs*^PlxnD1^* comprise subtypes with more specific laminar and projection patterns. We developed a novel combinatorial method to target PyN *^PlxnD1^* subtypes based on their birth order and anatomy (Fig. 6a). This method is based on the developmental principle that PyN birth time correlates with their laminar position, and the observation that the large majority of IT PyNs are generated from indirect neurogenesis through intermediate progenitors^36^.

We generated *PlxnD1-Flp;Tbr2-CreER;Ai65* compound mice by breeding the three alleles. In these mice, the constitutive *PlxnD1-Flp* allele marks the whole population (Fig. 6e) and the inducible *Tbr2-CreER* allele was used for birth dating PyNs*^PlxnD1^* (Fig 6a). Strikingly, TM induction at E13.5, 15.5 and 17.5 selectively labeled L5A, L3, and L2 PyNs *^PlxnD1^* (Fig. 6b-d). To reveal the projection pattern of these birth-dated PyNs, we bred the *PlxnD1-Flp;Tbr2-CreER;dual-tTA* compound allele mice for tTA-dependent viral injections. AAV-TRE3g-mRuby injection into SSp-bfd in E13.5- and 17.5-induced mice labeled distinct subtypes of PyNs*^PlxnD1^*, which differed in projection patterns. E13.5-born PyNs*^PlxnD1(E^*^13^*^.5)^* resided in L5A and projected ipsilaterally to multiple cortical areas, contralaterally to homotypic SSp-bfd cortex and heterotypic cortical areas such as the temporal association area (TEa), and bilaterally to the striatum (Fig. 6g-m, u). In contrast, E17.5-born PyNs*^PlxnD1(E17.5)^* resided in L2; although they also extended strong projections to ipsilateral cortical and striatal targets and to homotypic contralateral cortex, similar to PyNs*^PlxnD1(E13.5)^*, PyNs*^PlxnD1(E17.5)^* had minimal projections to heterotypic contralateral cortex and striatum (Fig. 6n-u).

Importantly, these two birth-dated PyNs*^PlxnD1^* subsets further differed in terms of their axon termination patterns within a cortical target area. Whereas PyNs*^PlxnD1(E13.5)^* axons terminated throughout the full thickness of L1 and L2/3, with few axon branches in L5A, PyNs*^PlxnD1(E17.5)^* axons terminated strongly in L2/3 and L5A, with few branches in L1 (Fig. 6l, m, s, t, u). This result revealed that, even within the same target areas, PyNs*^PlxnD1(E13.5)^* and PyNs*^PlxnD1(E17.5)^* may preferentially select different postsynaptic elements (e.g. cell types and/or subcellular compartments). The tight correlation between cell birth time, laminar location, projection target, and axon termination pattern suggests that the diversity of PyNs*^PlxnD1^* is unlikely to arise from stochastic processes but rather from the developmental program. This approach also demonstrates that it is feasible to target highly specific PyN subtypes by a strategic combination of developmental (lineage, birth order), molecular and anatomical attributes using just three alleles and one viral vector. The same approach can be combined with retrograde tracing approaches to further target PyN subtypes based on specific projection targets.

## DISCUSSION

A major principle of cell type organization in the cortex and across brain regions is their hierarchical taxonomy, consisting of top-level major classes that branch into progressively finer subpopulations and “atomic types”. An “ideal” genetic toolkit would have two essential features. The first is specificity at several levels of the hierarchy or granularity, from broader subclasses to finer types. The second is comprehensive coverage of many PyN subpopulations to enable systematic studies of cortical circuit organization. Here, we present genetic tools and strategies for targeting PyN subpopulations and tracking their developmental trajectories. Together with previous resources^15, 16, 69^, these driver lines provide substantial coverage of major PyN classes and subpopulations. The multiple inducible driver lines targeting progenitor types provide essential tools for fate mapping studies to explore the developmental mechanisms of PyN specification, differentiation, and circuit assembly. The combinatorial method that engages lineage, birth order, marker genes and anatomy allows highly specific targeting of PyN subtypes. This new set of PyN driver lines provides much improved specificity, robustness, reliability and coverage over existing resources. More importantly, the judicious choice and strategic design of driver lines based on developmental genetic mechanisms establish an experimental paradigm as well as a road map for dissecting the hierarchical organization of PyN types based on their inherent biology.

A fundamental difference between transgenic and gene KI approach to cell type targeting is the “nature” of regulatory mechanisms that drive tool-gene (e.g. CreER, Flp) expression. Transgene expression results from ectopic interactions of the transgenic promoter and enhancers with surrounding gene regulatory elements at a random genomic integration site^70^, giving rise to an arbitrary spatial and temporal “composite expression pattern” that often comprises an artificial mixture of cells. Desirable transgenic expression patterns are identified through screening multiple lines in which the same transgene is integrated at different genomic loci. Although useful once characterized, each transgenic expression pattern is unique (*i.e.* integration site-dependent) and often has no relationship to the transgene itself; such expression is not reproducible for designing intersection/subtraction patterns to target more restricted subpopulations. Furthermore, transgenic PyN driver lines often show unexplainable lack of expression in some cortical areas and unstable expression patterns. Thus existing PyN driver lines are inadequate for a systematic dissection of PyN types and progenitor types.

In contrast, the vast majority of KI lines precisely recapitulate the endogenous spatiotemporal and cellular gene expression patterns that reflect endogenous developmental processes, physiological mechanisms, and tissue organization. Thus, the usefulness of KI lines depends on the choice of targeted genes based on prior knowledge. Our effort has focused on key transcription factors and effector genes implicated in the specification, differentiation, and maintenance of PyN cell fate and phenotypes^3, 10, 18^. These KI driver lines thus enable investigators to parse biologically relevant PyN subpopulations through their inherent developmental, anatomical and physiological properties, i.e. “carving nature at its joints.” Further, as each driver line captures most if not all cells in each marker-defined subpopulation across cortical areas largely identical to mRNA in situ patterns, they enable investigators to study the global cortical areal network with cell type resolution. Importantly, the precision and reliability of KI drivers enable further combinatorial targeting of finer projection types through marker intersections and viral tools that incorporate anatomy and projection properties.

As most endogenous genes and transgenic constructs that show restricted expression to PyN subpopulations are often more broadly expressed at earlier developmental stages, many constitutive Cre lines are often not suited to take advantage of the valuable tool-gene reporter lines^69^ because Cre recombination patterns integrate across time and become non-specific to the intended PyN subpopulation in mature cortex. The temporal control of our inducible driver lines confers major advantages of specificity and flexibility in cell targeting, functional manipulation, and developmental fate mapping toward exploring circuit organization and assembly. Inducibility also allows investigators to control the density of labeling and manipulating PyN subpopulations, from dense coverage to single cell analysis. Single cell analysis provides the ultimate resolution to examine the variability and/or reliability of individual neurons within driver and anatomy defined PyN subpopulations^71^, and can be better achieved by inducible driver lines^72, 73^. Furthermore, temporal control allows gene manipulations (e.g. using conditional alleles) in specific subpopulations at different developmental stages to discover the cellular and molecular mechanisms of circuit development and function.

Several TFs used in this work may represent terminal selector genes involved in both the specification and maintenance of PyN cardinal identity (e.g. *Fezf2* for ET neurons, *Lhx2*, *Cux1/2* for IT neurons, *Tbr1* for CT neurons)^10, 17, 74^. Thus beyond experimental access to PyN subpopulations in mature cortex, these inducible TF driver lines represent powerful “functional fate mapping” tools for tracking the developmental trajectory of PyN types, from their specification to migration, differentiation, connectivity and circuit function. Unexpectedly, we found that *Fezf2* and *Tbr1*, which are considered class-defining TFs^10, 18^, do not perfectly parse the PT and CT classes, respectively. A small set of PyNs*^Fezf2^* co-expressing PlxnD1 in upper L5B and a set of PyNs*^Tbr1^* in L6 and/or L2/3 may project to contralateral cortex and striatum – an IT characteristic. As *Fezf2* and *Tbr1* are crucial fate-restricting TFs and each likely regulates a set of associated target genes, these results raise the intriguing possibility that the *Fezf2-* and *Tbr1-expressing* “IT-like” cells represent previously unrecognized novel or “intermediate” PyN types. This possibility can be examined by using intersectional strategies and single cell analysis to reveal their projection patterns, and the results may revise our understanding of how these master TFs shape PyN fate.

Importantly, a number of TFs used in this study continue to evolve across mammalian species from rodents to primates and human^75^ and might contribute to the development of human-specific traits. For example, *Cux1* is located in the human accelerated region (HAR) of the human genome^76^, and the regulation of *Cux1*, *Fezf2*, *Tbr1*, *Tbr2*, and *Foxp2* continues to evolve and diverge in primates and is implicated in several developmental disorders such as autism^35, 48, 65^. Therefore, these TF mouse driver lines may provide crucial handles to track the developmental trajectories of PyN subpopulations in the convoluted process of cortical circuit assembly, with implications in the evolutionary history of PyNs in human cortex. This will facilitate deciphering the genetic architecture of developmental disorders by linking gene mutations (e.g. with conditional alleles) to their impact on the developmental trajectory of specific neuron types and altered cortical circuit wiring and function.

Single cell transcriptomic analyses have identified dozens of marker gene combinations that define transcriptomic types^6^. Our study suggests that the strategic design of intersectional schemes, leveraging established and characterized driver lines, will substantially expand the scope and improve the specificity of PyN targeting. Recent advances in enhancer-based usage of AAV vectors show significant promise in achieving subpopulation restriction^77^. The intersection of enhancer AAVs with strategically designed driver lines may substantially increase the specificity, ease, and throughput of neuronal cell type access.

## Supporting information

Supplemental Video 1 Cux1;Ai14

Supplemental Video 2 Cux1;LSL-flp AAVfdTVAEGFP(SSp-bfd)

Supplemental Video 3 PlxnD1-CreER;Snap25-LSL-EGFP

Supplemental Video 4 PlxnD1-Flp;FSF-tdT

Supplemental Video 5 PlxnD1-CreER;LSL-flp AAVfdTVAEGFP(SSp-bfd)

Supplemental Video 6 Fezf2;Ai14

Supplemental Video 7 Fezf2;LSL-flp_AAVfdTVAEGFP(SSp-bfd)

Supplemental Video 8 Adcyap1;LSL-h2b-GFP

Supplemental Video 9 Adcyap1;LSL-flp AAVfdTVAEGFP(SSp-bfd)

Supplemental Video 10 Tcerg1l;LSL-h2b-GFP

Supplemental Video 11 Tcerg1l AAV-FLEX-EGFP(SSp-bfd)

Supplemental Video 12 Sema3E;Ai14

Supplemental Video 13 Tbr1;Ai14

Supplemental Video 14 Tbr1;LSL-flp AAVfdTVAEGFP(SSp-bfd)

Supplemental Video 15 Tle4;Snap25-LSL-EGFP

Supplemental Video 16 Tle4;LSL-flp AAVfdTVAEGFP(SSp-bfd)

Supplemental Video 17 Foxp2 SystemicAAV-CAG-FLEX-EGFP

Supplemental Video 18 Foxp2 AAV-CAG-FLEX-EGFP(SSp-bfd)

Supplementary Table 2 existing vs new

Supplementary Table 3 cell distribution traditional histology

Supplementary Table 4 cell distribution STPT

Supplementary Table 5 projections STPT

Supplementary Table 6 projections target values

Supplementary Table 7 projections traditional histology

Supplementary Table 8 Movie List

Supplementary Table 1 mouse lines

## EXTENDED DATA FIGURES

**Extended Data Fig. 1:**
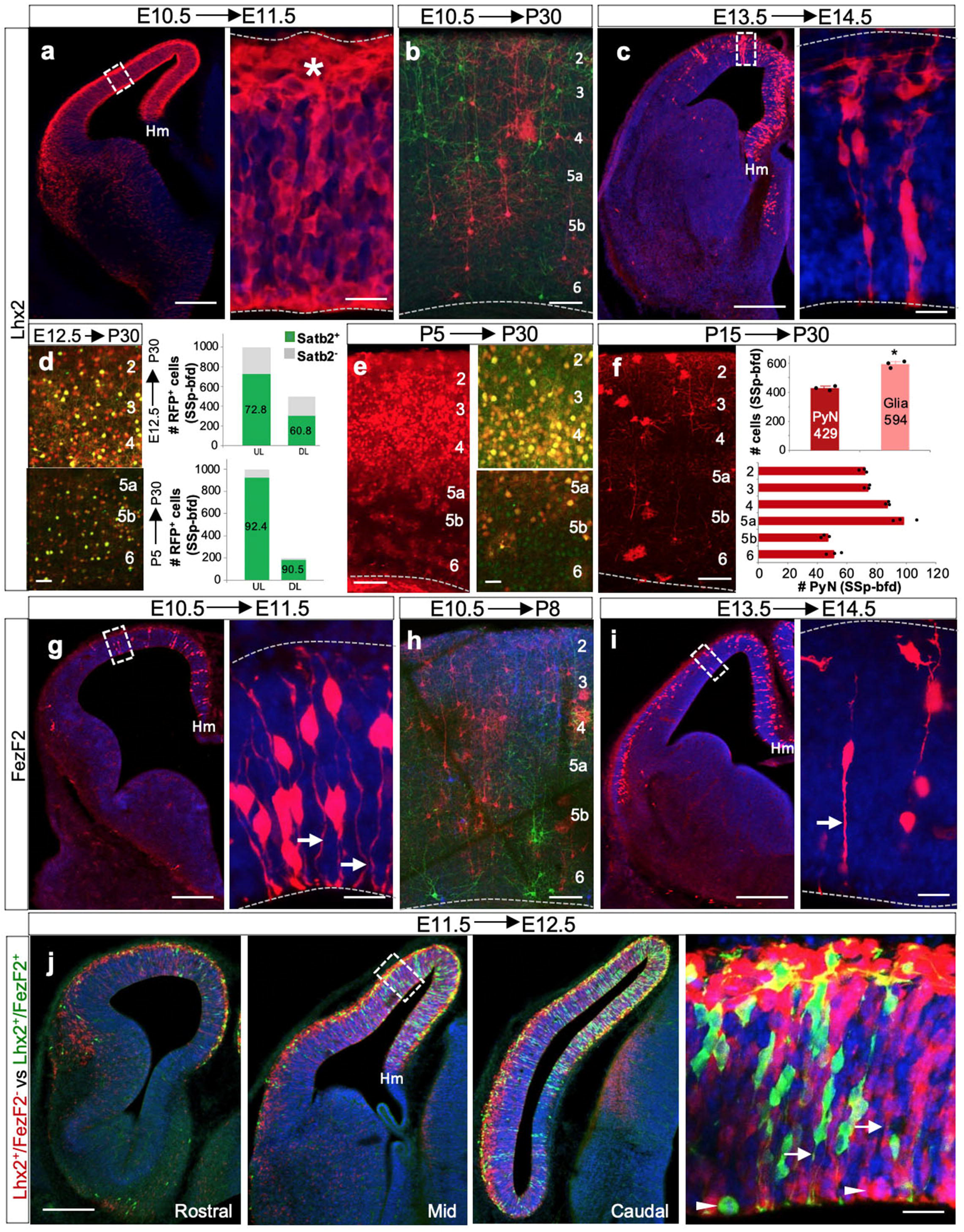
Fate mapping using *Lhx2* and *Fezf2* driver lines. a, A hemi-coronal view of E10.5 RGs*^Lhx2+^* densely labeled throughout the dorsal neuroepithelium by 24-hour pulse-chase in a different *Lhx2-CreER;Ai14* mouse as in Fig.2d. Hm, cortex-hem boundary. High-magnification views show a dense post-mitotic layer (asterisk) below pia (dashed lines). b, Another example of fate mapping from E10.5 RGs to mature cortex using *Lhx2-CreER;RGBow* mice (as in Fig.2e). E10.5 RGs*^Lhx2+^* are multipotent and generate PyNs across layers. c, same as in a except done at E13.5. High-magnification view shows cell clusters derived from presumably individual RGs indicative of the rapid proliferation potential of RGs*^Lhx2+^* at this time. d, Fate mapping E12.5 RGs*^Lhx2+^* to mature cortex labeled PyN progeny that are SATB2*^+^* (IT class, 66.8%) as well as SATB2*^-^* (non-IT class, 33.2%). Quantification: 72.8% of L2-4 (UL; 728 of 1000 cells) and 60.8% of L5-6 (DL; 304 of 500 cells; n=3 brains from 2 litters) PyNs are of IT-type. e, P5 TM induction in *Lhx2-CreER;Ai14* shows dense labeling of L2-4 PyNs and sparse labeling in L5/6 in P28 cortex. (Left) Most labeled PyNs are of the IT class expressing SATB2 (UL, 92.4% - 924 of 1000 cells; DL, 90.5% - 181 of 200 cells; n=3 brains from 2 litters). f, P15 TM induction in *Lhx2-CreER;Ai14*. (Upper) Quantification in SSp-bfd shows 58% cells are glia (Mean values for number of cells in SSp-bfd ± SEM. *P < 0.05 compared to PyN number, unpaired Student’s t test). (Lower) Labeled PyNs are distributed across cortical layers with more labeling in L5a and L4 (n=3 brains from 2 litters). g, At E10.5, RGs*^Fezf2+^* only be sparsely labeled by 24-hour pulse-chase using another *FezF2-CreER;Ai14* mouse (as in Fig.2m). Magnified view shows RGs with endfeet (arrow) but sparsely labeled post-mitotic neurons (arrowheads), compared to those of RGs*^Lhx2+^* at the same stage (Extended Data Fig1a). h, Fate mapping E10.5 RGs*^Fezf2+^* to P8 labeled PyNs across cortical layers in *Fezf2-CreER;RGBow* mice, indicating multi-potency of RGs*^Fezf2+^*. i, same as in g except done at E13.5; magnified view shows sparsely labeled RGs, in sharp contrast to the highly prolific RGs*^Lhx2+^* at the same embryonic time (Extended Data Fig1c). j, The presence of RGs*^Lhx2+Fezf2-^* (RFP) and RGs*^Lhx2+Fezf2+^*(GFP) at E11.5 throughout the cortical primordium is revealed by intersection/subtraction fate mapping with 24-hour pulse-chase in *Lhx2-CreER;Fezf2-Flp;IS* mice, schematized in Fig. 1c (Also see Fig. 2s-u). These progenitors distribute in a medial high-lateral low gradient along the dorsal neuroepithelium, ending at the cortex-hem boundary (Hm). Rostral, Mid and Caudal sectioning levels show a caudal high-rostral low distribution of RGs*^Lhx2+Fezf2-^* and RGs*^Lhx2+Fezf2+^*. Magnified view shows RGs*^Lhx2+Fezf2-^* and RGs*^Lhx2+Fezf2+^* at multiple cell cycle stages with endfeet (arrow) and dividing soma (arrowhead) at ventricle wall (dashed line). High magnification insets are not maximum intensity projections, and do not show the z-projection seen in the low mag image. Scale bars: 20μm in a,c,g,i,j; 100μm for all other panels.

**Extended Data Fig. 2:**
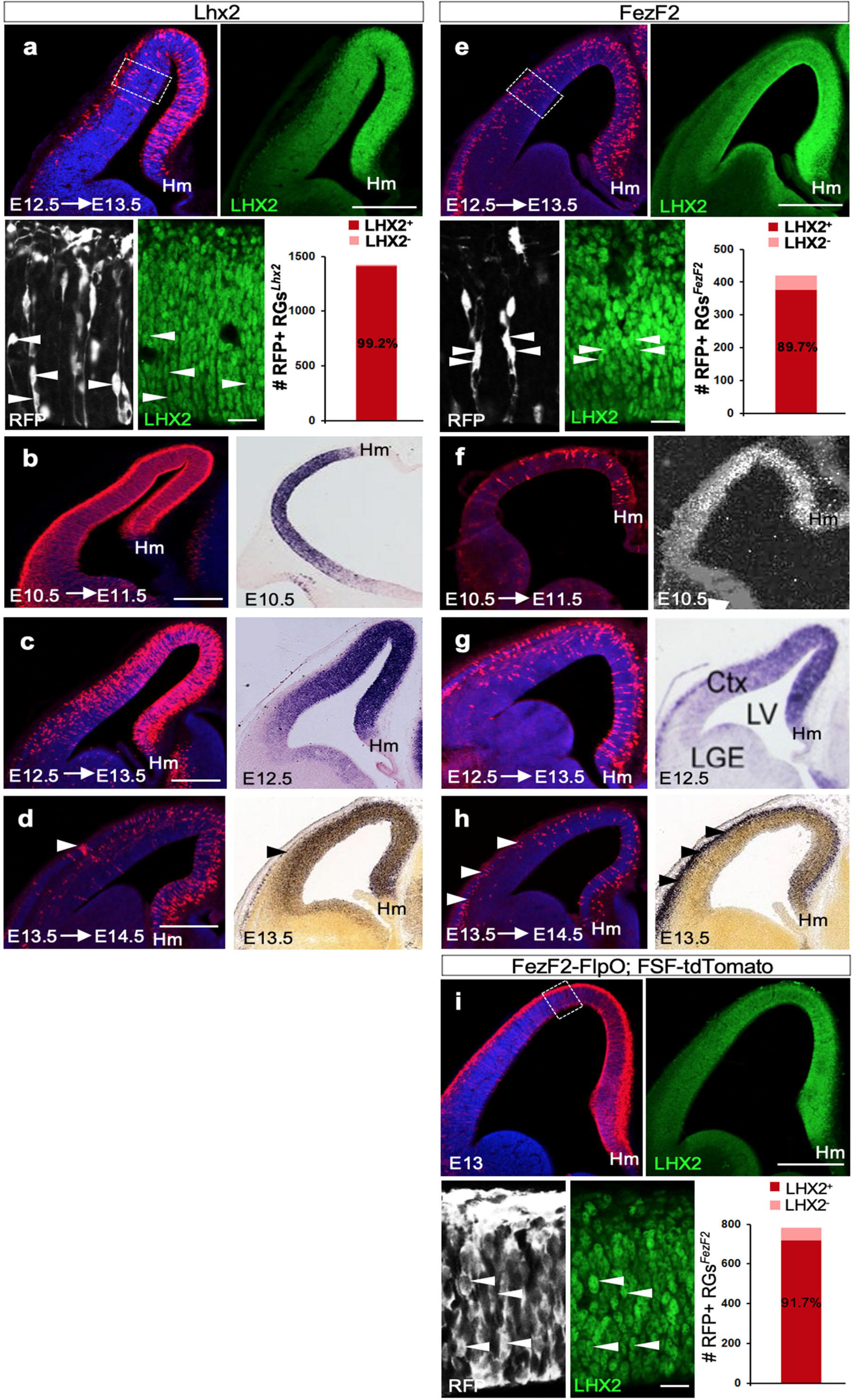
*Lhx2* and *Fezf2* driver lines precisely recapitulate endogenous developmental expression patterns. **a**, anti-LHX2 immunohistochemistry on a 24-hour pulse-chase in a E12.5 *Lhx2-CreER;Ai14* embryo reveals that 99.2% RGs*^Lhx2+^* express LHX2 (1410 of 1422 cells; n = 4 embryos from 2 litters). Colocalization seen in high magnification images (compare arrowheads; asterisk shows no colocalization). Note the medial^high^-lateral^low^ gradient. **b,** 24-hour pulse-chase in a E10.5 *Lhx2-CreER;Ai14* embryo densely labeled RGs throughout the dorsal neuroepithelium, ending medially at the cortex-hem boundary (Hm). This expression recapitulated *Lhx2* mRNA in-situ hybridization at E10.5 ^85^. **c**, Same experiment as in **b** done at E12.5. The medial^high^-lateral^low^ gradient of RGs*^Lhx2+^* is highly similar to *Lhx2* mRNA expression ^27^. **d**, Same experiment as in **b** done at E13.5. Note the reduction in RGs*^Lhx2+^* and concomitant increase in post-mitotic cells (arrowheads), in both the *Lhx2* driver line and mRNA expression (ISH Data: Allen Brain Atlas: Developing Mouse Brain). **e**, anti-LHX2 immunohistochemistry on a 24-hour pulse-chase in a E12.5 *FezF2-CreER;Ai14* embryo reveals that 89.7% RGs*^FezF2+^* express LHX2 (376 of 419 cells; n =4 embryos from 2 litters). Colocalization seen in high magnification images (compare arrowheads) **f**, 24-hour pulse-chase in a E10.5 *Fezf2-CreER;Ai14* embryo. The spatial extent of RGs*^Fef2+^* is restricted compared to RGs*^Lhx2+^* at this stage. This sparse labeling recapitulated *FezF2* mRNA expression at E10.5 ^27^. **g**, Same experiment as in **f** done at E12.5. The medial^high^-lateral^low^ distribution gradient of RGs*^Fezf2+^* is also seen with *FezF2* expression at this stage. Note RGs*^Fezf2+^* are sparsely labeled compared to RGs*^Lhx2+^* using in-situ hybridization at E12.5 ^86^. **h**, Same experiment as in **f** done at E13.5. The drastic decrease of *FezF2* expression in RGs^FezF*2+*^ accompanied by an increase in post-mitotic PyNs (arrowheads) is comparable in both *FezF2-CreER; Ai14* and *Fezf2* in-situ data (ISH Data: Allen Brain Atlas: Developing Mouse Brain). **i,** anti-LHX2 immunohistochemistry on E13 *Fezf2-FlpO;FSF-tdTomato* embryo reveals that 91.7% RGs*^Fezf2+^* express LHX2 (718 of 783 cells; n = 4 embryos from 2 litters). Also compare with **e**. Colocalization seen in high magnification images (compare arrowheads; asterisk shows no colocalization). Note the medial^high^-lateral^low^ gradient. Scale bars for low mag images = 100μm in **a-i**. Scale bars for high mag images = 20μm in **a,e,i**.

**Extended Data Fig. 3:**
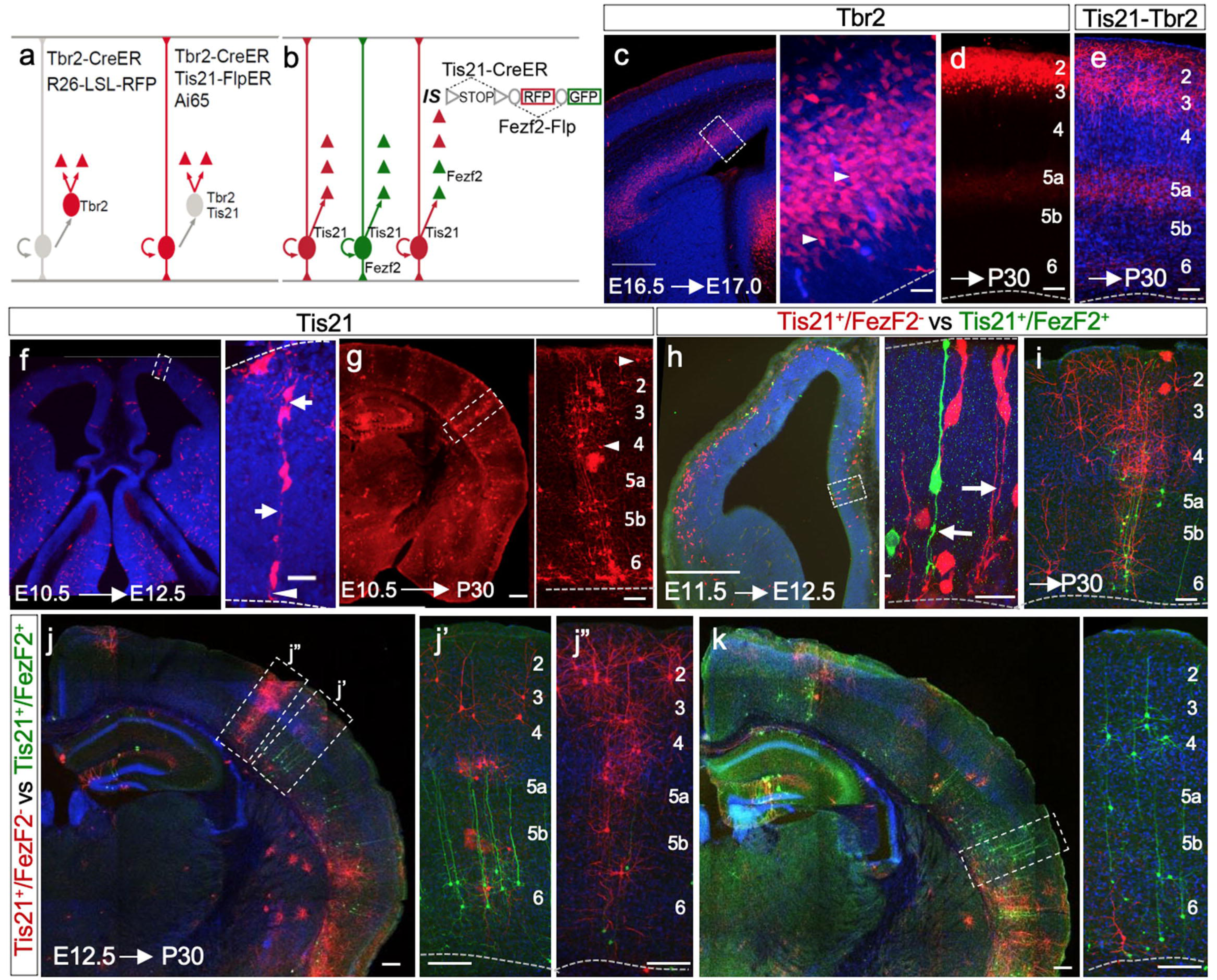
Fate mapping of neurogenic and intermediate progenitors. **a,** Fate mapping strategy for intermediate progenitors (IPs) and indirect neurogenesis, used in **c-e**. *Tbr2*-CreER labels an IP and all of its progeny with a fluorescent marker when combined with Ai14 (Left). The intersection of *Tis21-CreER* and *Tbr2-FlpER* specifically targets neurogenic IPs when combined with the Ai65 intersectional reporter (Right). **b**, Simultaneous fate mapping of different molecularly defined neurogenic RGs using an intersection/subtraction reporter (*IS*) combined with *Tis21-CreER* and *Fezf2-Flp* drivers. This scheme is used in **h-k**. In RGs*^Tis21+Fezf2-^*, Cre activates RFP expression only. In RGs*^Tis21+Fezf2+^*, Cre and Flp recombinations remove the RFP cassette and activate GFP expression. At a later stage when *Fezf2* is only expressed in postmitotic deep layer PyNs, *Tis21-CreER* in RGs activates RFP expression in all of its progeny, but RFP is then switched to GFP only in *Fezf2^+^* PyNs expressing Flp. **c**, E16.5 IPs densely labeled by 12-hour pulse-chase in *Tbr2-CreER;Ai14* mice; magnified view shows IP somata (arrowhead) away from the lateral ventricle (dashed line) lacking radial fibers and endfeet. **d**, Fate mapping E16.5 IPs in *Tbr2-CreER;Ai14* mice labels PyNs in L2-3 cortex at P28. **e**, Intersectional fate mapping of neurogenic IPs at E16.5 in *Tis21-CreER;Tbr2-FlpER;Ai65* mice, as depicted in **a**, labeled L2-3 PyN progeny in P28 cortex. **f,** 48-hr pulse-chase in E10.5 *Tis21-CreER;Ai14* embryo labels *Tis21^+^* neurogenic progenitors (nRGs) and their postmitotic progeny throughout the neural tube, including dorsal pallium (high magnification). Self-renewing RGs are identified by their endfeet at the ventricular surface (arrowheads) and radial fibers (arrow). **g**, Fate-mapping of E10.5 nRGs to mature cortex reveals PyNs are distributed throughout cortical layers. Note that multipolar GABAergic interneurons (some in layer 1) derived from subpallium nRGs are also labeled (arrowheads). **h**, The presence of nRGs*^Fezf2-^* and nRGs*^Fezf2+^* at E11.5 is revealed by intersection/subtraction fate mapping with 24-hour pulse-chase in *Tis21-CreER;Fezf2-Flp;IS* mice, schematized in **b**; magnified view shows RFP-labeled nRGs*^Fezf2-^* and GFP-labeled nRGs*^Fezf2+^*. **i**, Fate mapping E11.5 nRGs using *Tis21-CreER;Fezf2-Flp;IS* mice. The mixed RFP and GFP clone is likely derived from a nRG*^Fezf2-^*, which activated RFP expression in all progeny and GFP expression was then switched on only in *Fezf2^+^* postmitotic deep layer PyNs expressing Flp. **j-k**, More examples of differential fate mapping of nRGs*^Fezf2-^* and nRGs*^Fezf2+^* from E12.5 to the mature cortex using the scheme in **b**. The majority of clones consist of mixed RFP and GFP PyNs (j’), and rarely RFP-only (j”) or GFP-only PyNs (k). RFP-only clones (1 of 29) likely derive from nRG*^Fezf2-^* whose progeny were all *Fezf2*^-^ (j”). GFP-only clones (2 of 29) are derived from nRGs*^Fezf2+^*, suggesting multipotency of RGs*^Fezf2+^* (k). Mixed RFP/GFP clones are most prominent and likely result from Cre activation of RFP in nRGs*^Fezf2-^* and subsequent Flp activation of GFP in *Fezf2^+^* L5/6 postmitotic PyNs (i,j’). Scale bars: 500μm in **d,j,k**; 100μm in **d** (high mag)**,e,f,g,h,i,j’,j”,k** (high mag); 20μm in **c,e,h**.

**Extended Data Fig. 4:**
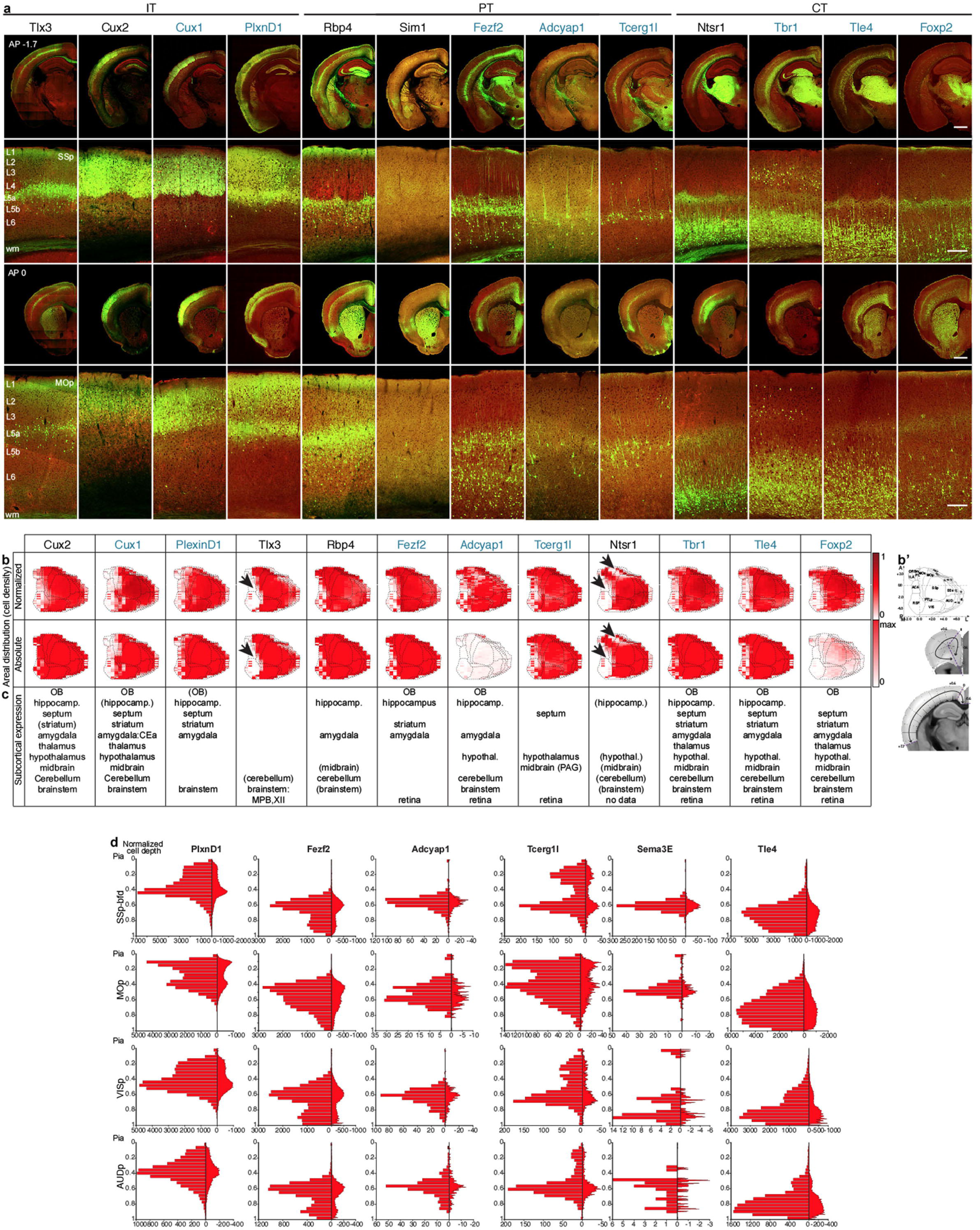
Comparison with existing driver lines in terms of areal and laminar patterns. **a,** Side-by-side comparison of Cre recombination patterns from mouse driver lines characterized in this study (blue font) and existing driver lines (black font) visualized through reporter expression (green; background autofluorescence in red), grouped according to IT, PT and CT projection classes. First row: coronal hemisections at Bregma −1.7 mm. Second row: Image panel showing cortical depth detailing cell body distribution pattern of PyN subpopulations within SSp-bfd taken from the hemisection above at level Bregma −1.7 mm. Third row: coronal hemisections at Bregma 0 mm. Image panel showing cortical depth detailing cell body distribution pattern of PyN subpopulations within MOp taken from the hemisection above at level Bregma 0 mm. **b,** Cortex-wide distribution patterns of PyN subpopulations viewed as cortical flatmaps in a side-by-side comparison of newly generate (blue font) and existing driver lines (black font): first row, normalized for each dataset’s total number of cells detected; second row, absolute scale per flatmap grid area, with maximum number of cells for any PyN subpopulation. **b’** shows cortical flatmapping coordinate space and two exemplary coronal hemisections describing the demarcations used to generate the cortical grid for flatmapping ^81^. **c,** Overview of brain-wide cell body distribution patterns for each driver line. This table provides an overall impression of the recombination patterns in major adult brain regions in selected lines. **d,** Histograms showing normalized laminar distribution for six genetically targeted PyN subpopulations by cortical area. Brain-wide cortical depth quantification was performed based on cell detection by convolutional networks from GeneX-CreER driver lines crossed to Ai14 (R26-LSL-tdTomato), R26-LSL-h2b-GFP or Snap25-LSL-EGFP reporters and induced at the ages specified in Figures 1 and 3. The normalized cortical depth (0-1) was divided into 24 bins for the left histogram and 124 bins for the right plot in each panel. Abbreviations: SSp, Primary somatosensory cortex; MOp, Primary motor cortex; VISp, Primary visual cortex; and AUDp, Primary auditory cortex; AP, antero-posterior coordinate relative to Bregma. Scale bars: Last panel of first and third rows applies to all hemisections, 1mm; last panel of second and fourth rows applies to all cortical depth image panels, 200μm.

**Extended Data Fig. 5:**
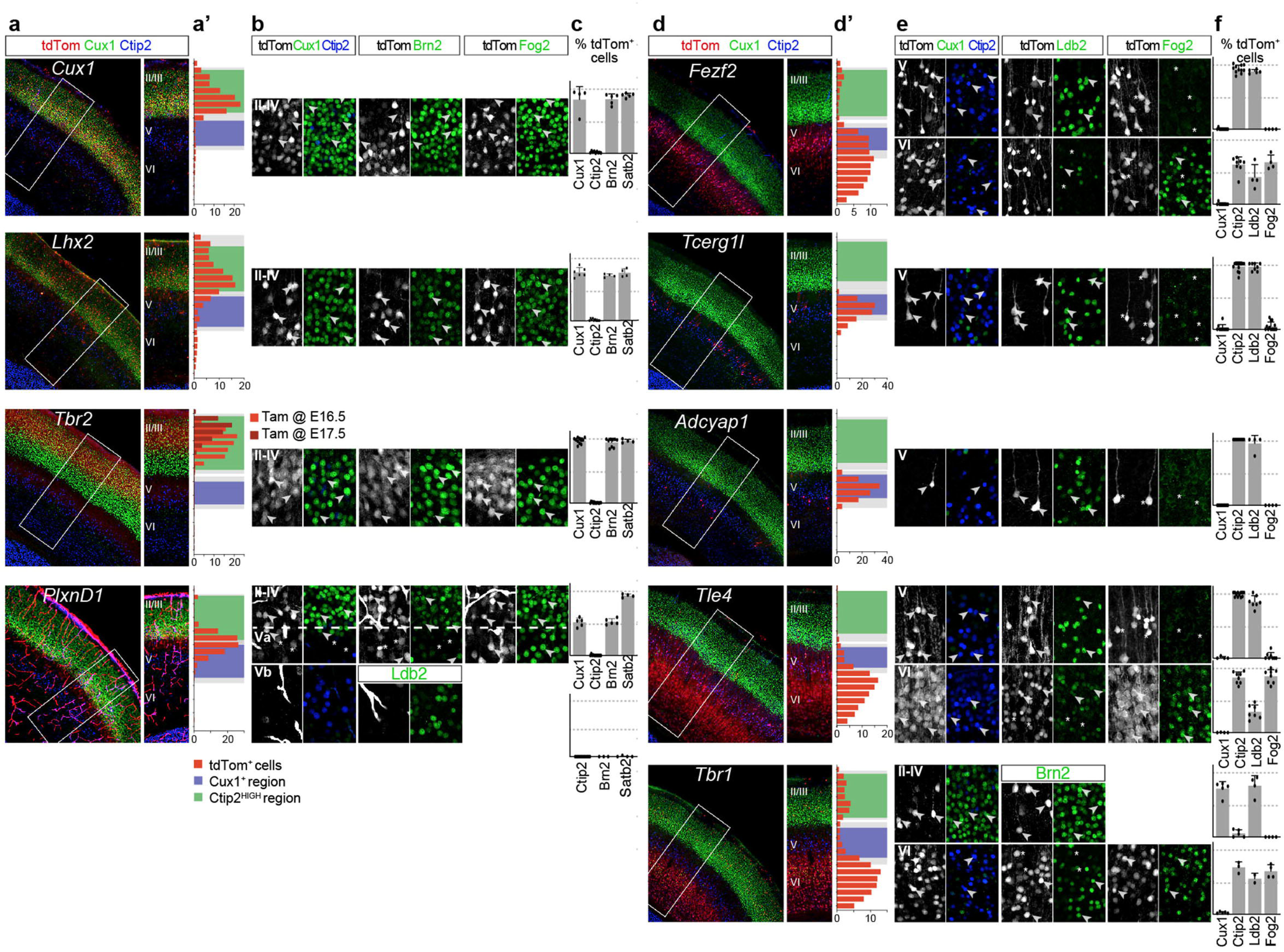
Molecular validation and characterization of PyN driver lines. **a,d,** Low magnification images of sections of somatosensory cortex at P7 stained with antibodies against CTIP2, CUX1, and tdTomato from *PyN*-*CreER;Ai14* mice induced with tamoxifen. Inset to the right shows markers and Tomato^+^ cell distributions across layers. *Fezf2*-, *Tcerg1l*-, *Adcyap*- and *Tle4*-CreER;Ai14 were induced at E16.5 and collected at P7. *Tbr1*-, *Cux1*- and *Plxnd1*-CreER;Ai14 were induced at P4 and collected at P7. *Tbr2*-CreER;Ai14 was induced at E16.5 (not shown) or E17.5 and colected at P7. *Lhx2*-CreER;Ai14 was induced at P3 and collected at P7. **a’,d’,** Histograms showing the radial distribution of Tomato^+^ cells in the cortical plate, in the region corresponding to the somatosensory cortex. Briefly, sections stained with CUX1 and CTIP2 were used and the position of Tomato^+^ cells, relative to the thickness of the cortex was recorded, as well as the limits of the areas occupied by CUX1^+^ or CTIP2^HIGH^ cells, shown in green (layers 2-4) and blue (layer 5b) bars, respectively (average relative values for the same sections, gray shading corresponds to 1 SD). For *Tbr2*-CreER;Ai14, animals induced at E16.5 or E17.5 were quantified separately, showing the later induction (darker red) results in more superficial labelling. Quantifications were made from 4-10 sections from 2-3 different mice for each line. **b,e,** Magnification of Tomato labelled cells in sections co-stained against CUX1 and CTIP2, LDB2 (enriched in PT), FOG2 (expressed in CT), BRN2, or SATB2 (expressed in IT). Arrowheads show double positive cells, while asterisks show Tomato^+^ cells not expressing the marker. **c,f,** Percentage of Tomato^+^ cells stained with each antibody. Quantifications were done in equivalent areas (320m by 320 m) within the somatosensory cortex centered in the specified layers. Each dot is an area from a different section, for which the percentage of double positive cells was calculated. Bars are mean+SD. Quantifications were made from 4-8 sections from 2-3 different animals for each line. *Tbr2*-CreER;Ai14 labelled CUX1^+^, SATB2^+^, BRN2^+^ IT in the most superficial layers 2-3, irrespective of their induction time. *Lhx2*-CreER;Ai14 and *Cux1*-CreER;Ai14 labelled CUX1^+^, SATB2^+^, BRN2^+^ IT deeper in layers 2-3. *Plxnd1*-CreER;Ai14 labelled SATB2^+^, BRN2^+^ cells in layer 5a, as well as CUX1^+^, SATB2^+^, BRN2^+^ cells in layer 4. No cells were found in layer 5b. *Tcerg1l*-CreER;Ai14 and *Adcyap*-CreER;Ai14 labelled sparse LDB2^+^, CTIP2^+^ PT in Layer 5. *Fezf2*-CreER;Ai14 extensively labelled PT in layer 5 that were LDB2^+^, CTIP2^+^, as well as some CT in layer 6 expressing CTIP2 and FOG2. *Tle4*-CreER;Ai14 and *Tbr1*-CreER;Ai14 labelled CT expressing FOG2 and CTIP2 (and lower levels of LDB2) in layer 6. *Tle4*-CreER;Ai14 also labelled some LDB2^+^, CTIP2^+^ cells in layer 5 (PT), while *Tbr1*-CreER;Ai14 also labelled some CUX1^+^, BRN2^+^ IT in layer 2/3.

**Extended Data Fig. 6:**
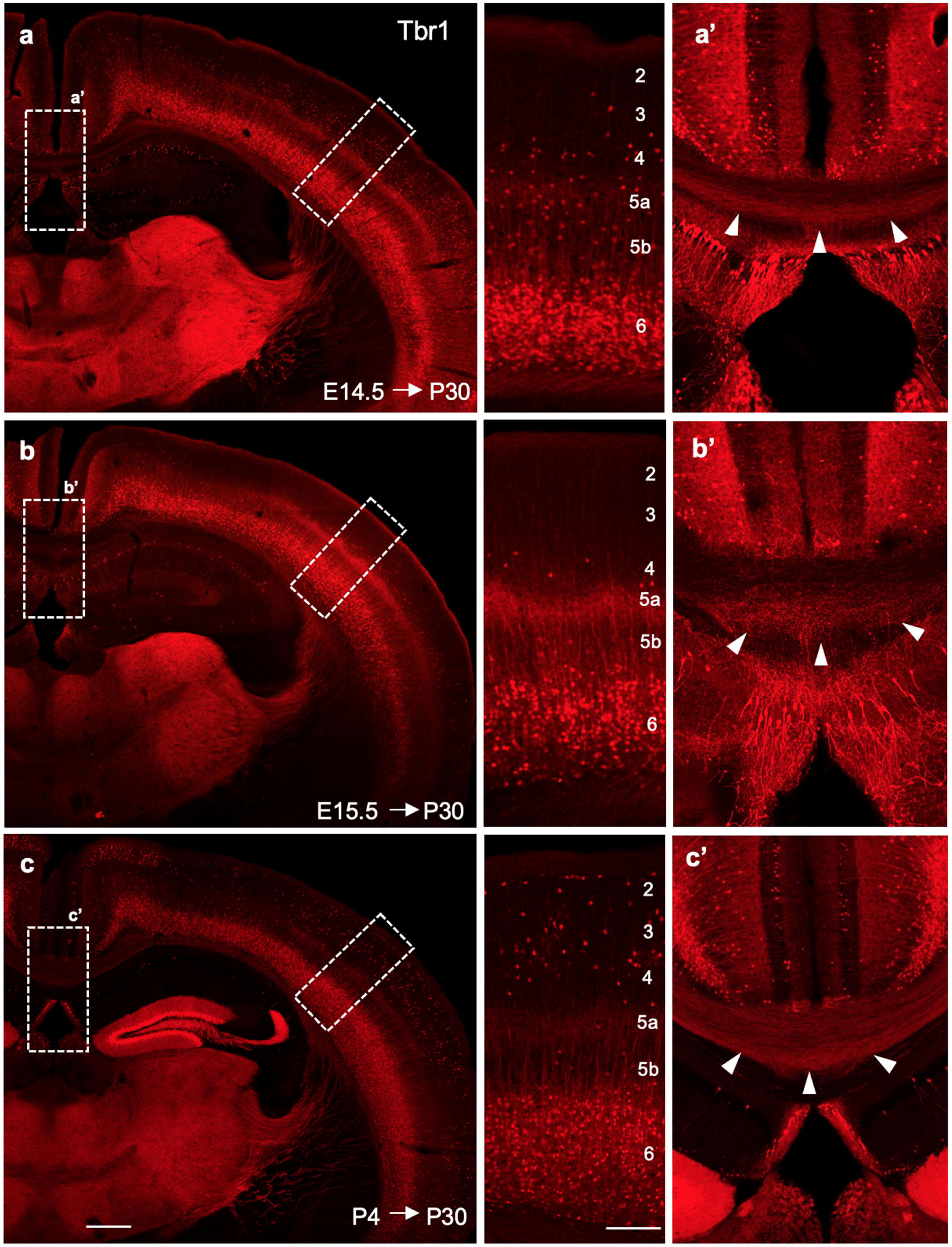
*Tbr1* is largely restricted to L6 CT PyNs and does not broadly label all newly born post-mitotic cortical PyNs. **a,** Fate mapping of PyNs using *Tbr1*-CreER; Ai14 (PyNs*^Tbr1^*). TM induction at E14.5 densely labels L6 CT cells with minor labeling of cells in L3-5. **b,** TM induction at E15.5 labels L6 CT cells. **c**, TM induction at P4 labels L6 CT cells and also a subset of L2/3 cells. A subset of adult PyNs*^Tbr1^* labeled from E14.5, E15.5 and P4 induction project to the contralateral cortex via the corpus callosum (arrowheads, **a’, b’, c’**). Scale bars for low mag = 500μm, and for high mag = 100μm.

**Extended Data Fig. 7:**
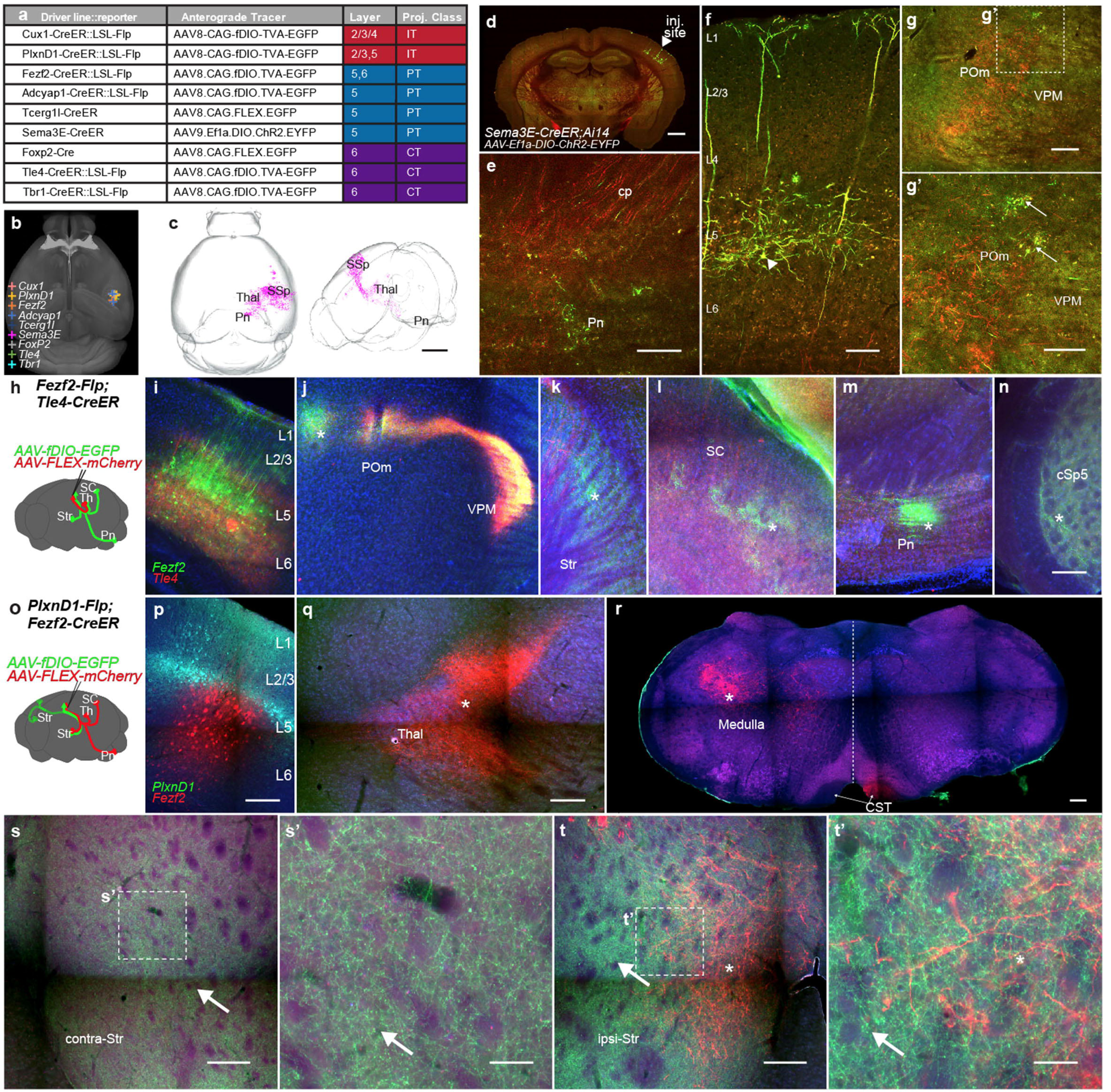
Anterograde tracing of axon projections from one or two PyN populations using single or double allele driver mice, respectively. **a,** Summary table of driver lines and viral vectors used for anterograde tracing from PyNs in primary somatosensory cortex, related to Fig. 4. **b,** Virus injection centroid coordinates across single driver experiments in CCFv3 space on a dorsal whole-brain view. **c,** 3D rendering of PyNs^Sema3E^ projection pattern from STP imaging: dorsal view (left) and parasagittal view (right). **d,** Coronal section containing injection site in a *Sema3E*-*CreER;Ai14* mouse. **e,** PyNs^Sema3E^ project via the cp to Pn. PyNs^Sema3E^ infected with AAV-DIO-EYFP express EYFP/tdTomato; all other PyNs^Sema3E^ express tdTomato by TM induction. **f,** PyNs^Sema3E^ at injection site, showing somata in layer 5 with slender tufted apical dendrites. **g,** PyNs^Sema3E^ axons in thalamus with large boutons in POm (**g’**). **h-t,** Simultaneous anterograde tracing from two driver allele-defined PyN populations. **h,** Schematic showing simultaneous anterograde tracing from PyNs targeted by *Fezf2*-Flp (green) and *Tle4*-CreER (red) with co-injection of Flp- and Cre-dependent AAVs expressing EGFP and mCherry, respectively (**i-n**). **i,** PyNs^Fezf2^ and PyNs^Tle4^ at the injection site occupying mainly L5B and L6, respectively. **j,** PyN^Fezf2^ and PyN^Tle4^ projection patterns converge in primary thalamus, VPM, while PyNs^Fezf2^ collaterals (asterisk) extend medially to higher order thalamic nuclei. **k,** PyNs^Fezf2^ (green) extend axon collaterals in Str, while PyNs^Tle4^ (red) pass through en route to thalamus. **l-n,** PyNs^Fezf2^ but not PyNs^Tle4^ project to multiple other corticofugal targets, including SC, Pn and cSp5. **o,** Schematic showing simultaneous anterograde tracing from PyNs targeted by *PlxnD1-Flp* (green) and *Fezf2-CreER* (red) with co-injection of Flp- and Cre-dependent AAVs expressing EGFP and mCherry, respectively (**p-t**). **p,** PyNs^PlxnD1^ and PyNs^Fezf2^ at the injection site in motor cortex occupying mainly L5A and L5B/L6, respectively. **q-r,** PyNs^Fezf2^ but not PyNs^PlxnD1^ project to Thal (**q**) and medulla (**r**). **s-t,** PyNs^PlxnD1^ and PyNs^Fezf2^ project to ipsilateral Str with overlapping terminals (**t**), while PyNs^PlxnD1^ but not PyNs^Fezf2^ project to contralateral Str **(s)**. Asterisks indicate PyN^Fezf2^ collaterals and arrows indicate PyN^PlxnD1^ collaterals. Scale bars: **c,** 2 mm; **d,** 1 mm; **e-g,** 200 μm; **g’,** 100μm; **n** (applies to i-n), 200μm; **p-t,** 200μm; **s’ & t’,** 100μm. Abbreviations: cp, cerebral peduncle; Pn, pons; Thal, thalamus; POm, posteromedial complex of the thalamus; VPM, ventral posteromedial nucleus; Str, striatum; SC, superior colliculus; cSp5, contralateral spinal trigeminal nucleus; CST, corticospinal tract.

**Extended Data Fig. 8:**
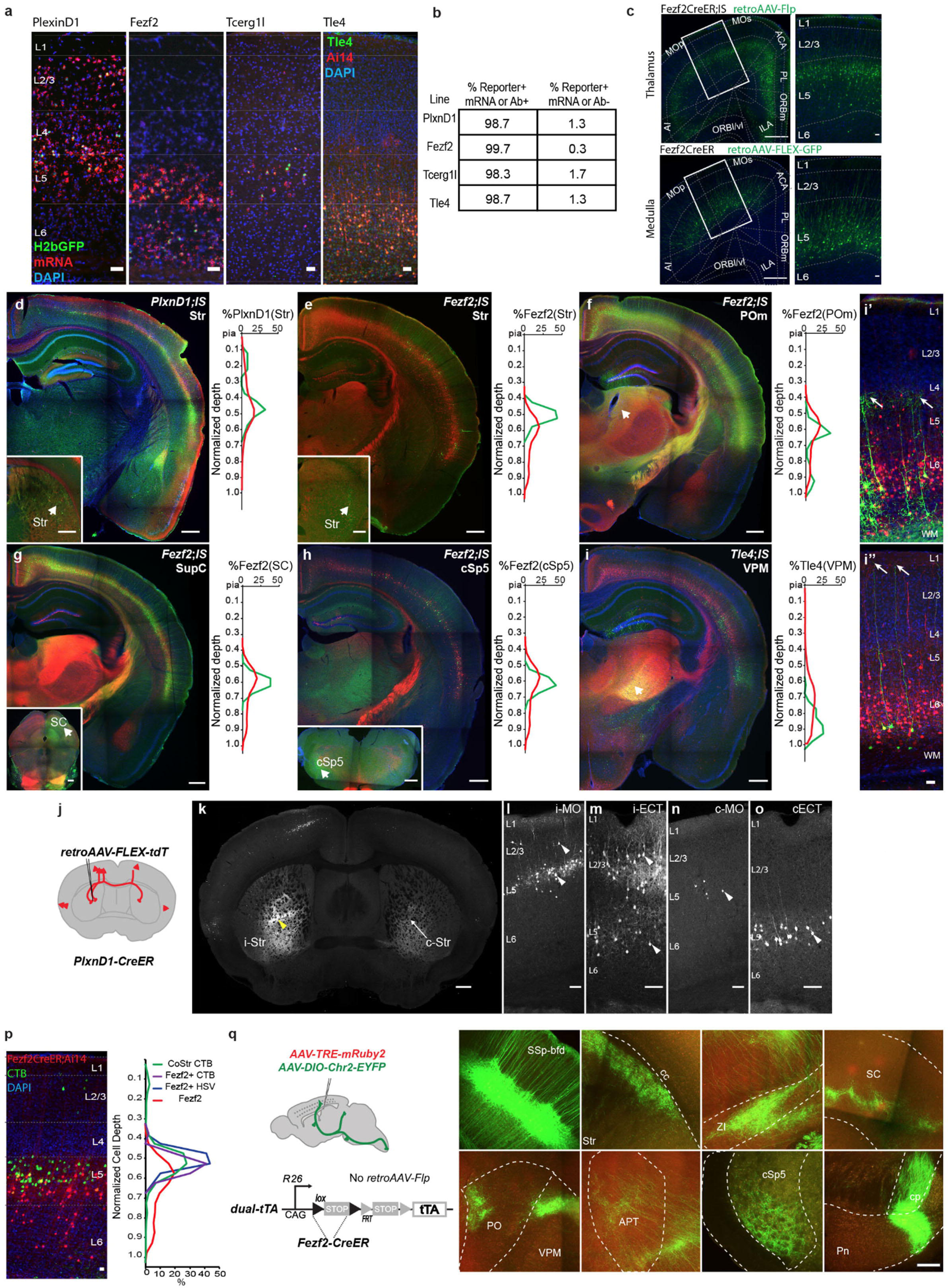
Retrograde and combinatorial targeting of PyN subpopulations. **a,** Validation of four PyN driver lines by fluorescent in situ hybridization (*PlxnD1, Fezf2, Tcerg1l* and antibody (TLE4). *PlxnD1-, Fezf2-, Tcerg1l-Cre* drivers were bred with a Rosa26-loxpSTOPloxp-H2bGFP reporter. H2bGFP signals co-localized with mRNA in situ signals of *PlxnD1-, Fezf2-, Tcerg1l*. In *Tle4-Cre;Ai14* mice, RFP signals co-localilzed with immunofluorescence of the TLE4 antibody. Scale bars are 50 um. **b,** Quantification of the co-localization of the Cre recombination signals with endogenous mRNA (*PlxnD1, Fezf2, Tcerg1l*) or protein (TLE4) signals. **c,** Another experiment related to Fig. 5e, showing that PyNs^Fezf2^ retrogradely labeled from thalamus and medulla are distributed in the upper or lower L5B, respectively, in the motor cortex. In a *Fezf2-CreER;IS* mouse (upper panels), retroAAV-Flp was injected in thalamus. In a *Fezf2-CreER* mouse (lower panels), retroAAV-Flex-GFP was injected into the medulla. Scale bars are 500 um (left column) and 50 um (right column). **d-i,** Representative hemi-sections containing the SS-bfd showing the retrograde labeling patterns PyNs^PlxnD1^, PyNs^Fezfe^, PyNs^Tle4^ by retroAAV-Flp injections at subcortical targets (arrow heads in insets or panels) in a *PyN-CreER;IS* mice. In each panel, the whole population in each cortex was labeled by RFP, whereas the retrograde labeled subset express GFP. *Fezf2* hemisections correspond to image panels in Fig. 5g. Corresponding cortical soma depth distribution is shown to the right of each image panel (n = 2 for each target). PyNs^Tle4^ project to the ventral posteromedial nucleus of thalamus (VPM) and consist of two subpopulations with apical dendrites in L4/5 (**i’**) and L1 (**i’’**), respectively, indicated by arrows. **j,** Retrograde targeting of striatum-projecting PyNs^PlxnD1^ by injection of retroAAV-FLEX-tdTomato in striatum. **k**, Coronal section displays injection site (arrowhead) and collaterals of retrogradely labeled PyNs^PlxnD1^ in contralateral striatum (arrow). **l-o,** Laminar pattern of retrograde labeled PyNs^PlxnD1^ reveal what while L5A PyNs^PlxnD1^ project to both ipsi- and contralateral side (**l, m, n, o**), L2/3 PyNs^PlxnD1^ mainly project to the ipsilateral striatum (**l, m**). **p,** Absence of tropism effect on retrograde PyNs^Fezf2^ labeling. Comparison between retrograde CTB injection in striatum of *Fezf2-CreER;Ai14* (N=3, coronal section shown on left, scale bar is 50 um) and HSV-Flp injection in striatum of *Fezf2-CreER;IS* mice (images in Fig. 5g) to label striatum-projecting PyNs^Fezf2^ (CoStr) in SSp. Depth distributions compare total CTB-labeled PyNs (green), CTB-(magenta) and HSV- (blue) labeled PyNs^Fezf2^, and the whole population of PyNs^Fezf2^ labeled in *Fezf2-CreER;Ai14* in SSp (red). Both CTB and HSV labeled more superficial L5 biased PyNs^Fezf2^, showing identical laminar labeling pattern between the labeling methods. **q,** Schematic of control experiment for use of Cre- and Flp-dependent dualtTA for target-defined axon projection mapping of PyNs^Fezf2^ (‘triple trigger’, related to Fig 5i**,j**). Co-injection of a Cre-dependent AAV-DIO-ChR2-EYFP (green, positive control) and tTA-activated AAV-pHB-TRE-mRuby2 (red, negative control), followed by TM induction, in absence of Flp probes dependence of reporter on both Cre and Flp recombination. Example images of injection site (SSp-bfd) and axon projection targets of several indicated ipsi- and contra-lateral sites display EGFP+ PyNs^Fezf2^ axons from AAV dependent on Cre alone, but no mRuby2+ axons, demonstrating the dependence of triple trigger strategy (Fig. 5f) on intersection of Cre and Flp. Scale bar in: a, 50μm; c, 500μm, 50μm; d-i, 500μm; i’-i’’, 50μm; k, 500μm; l-o, 100μm; p, 50μm; q, 200μm.

**Extended Data Fig. 9:**
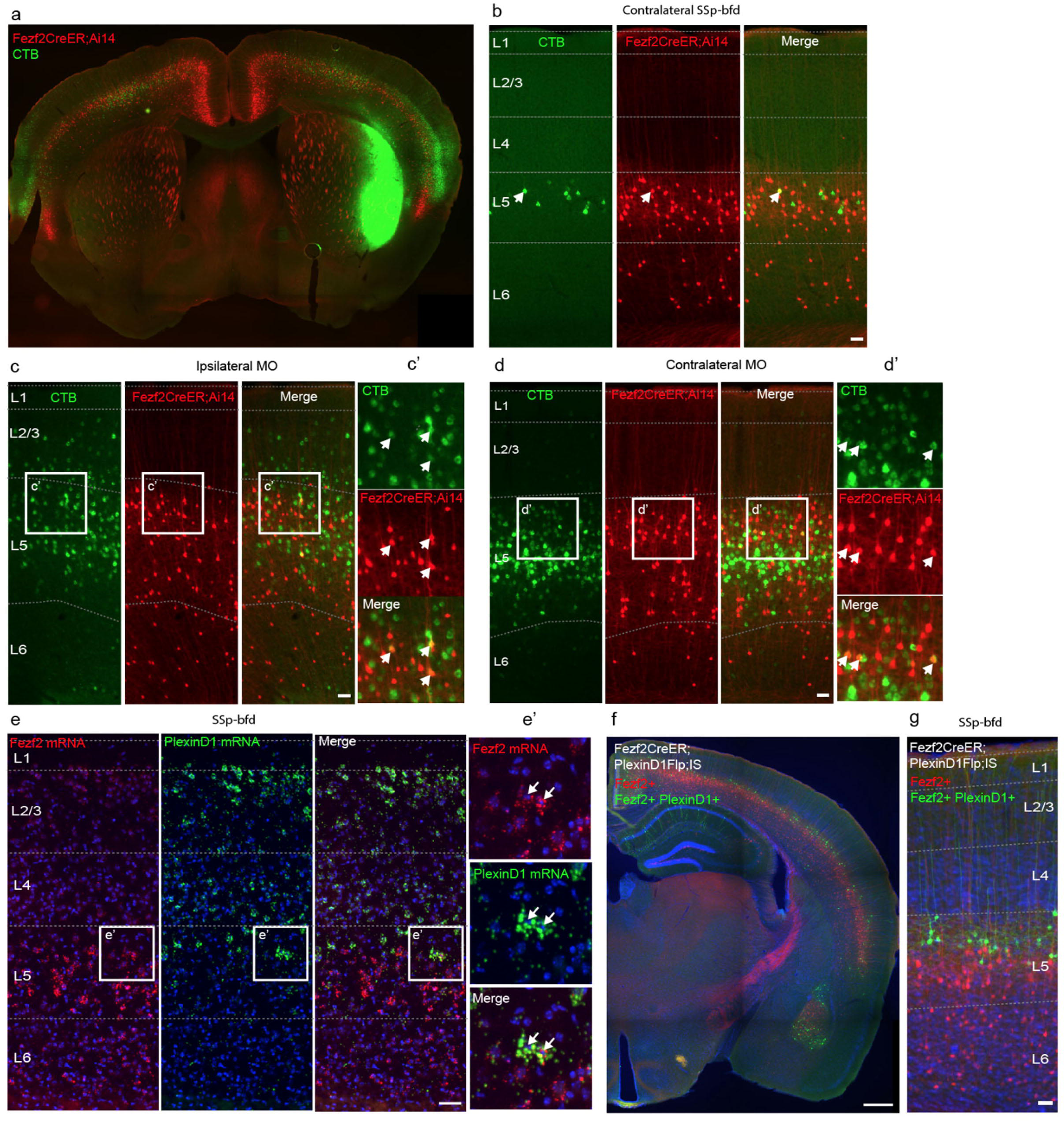
A subset of PyNs^Fezf2^ manifest IT features. **a-d**. Retrograde injection of CTB in the lateral striatum labeled a set of contralateral PyNs^Fezf2^ at the top of L5B in SSp-bfd (**a,b**), and ipsilateral (**c, c’**) and contralateral (**d, d’**) MO. **e**, *Fezf2* and *PlxnD1* mRNAs are co-expressed (**e’,** arrows) in a subset of L5 PyNs in the L5B and L5A border region. **f**, **g,** Intersection/subtraction mapping in *Fezf2-CreER;PlxnD1-Flp;IS* mice revealed that a small set of PyNs^Fezf2/PlxnD1^ (GFP) were located at the top of L5B among most other PyNs^Fezf2^ (RFP). Magnified insets are from the boxed regions in each panel. DAPI (blue) counterstain in e-g. Scale bars: a, f, 500μm; b, c, d, g, 50μm.

## SUPPLEMENTARY TABLES

**Extended Data Table 1** Newly generated mouse driver lines targeting cortical PyNs.

**Extended Data Table 2** Comparison of new and existing driver lines.

**Extended Data Table 3** Summary of cell distribution experiments acquired by traditional histology.

**Extended Data Table 4** Summary of cell distribution datasets acquired by STPT.

**Extended Data Table 5** Summary of projection datasets acquired by STPT.

**Extended Data Table 6** Full list of SSp-bfd axon targets and values measured from automated detection. Values report number pixels reported per brain structure per dataset on right and left hemisphere.

**Extended Data Table 7** Summary of projection datasets acquired by traditional histology.

**Extended Data Table 8** List of movies acquired by STPT

## LIST OF ABBREVIATIONS

AAV: adeno-associated virus
ACA: anterior cingulate area
AI: Agranular Insula
AMY: amygdala
AP: antero-posterior coordinate relative to Bregma
APT: anterior pretectal nucleus
AUDp: Primary auditory cortex
BIL: BICCN Brain Image Library
BLA: basolateral amygdala
BS: brainstem
Cb: cerebellum
cc: corpus callosum
c-ECT: contralateral ectorhinal cortex
c-MO: contralateral motor cortex
CNU: cerebral nuclei
cp: cerebral peduncle
CPN: callosal projection IT PyN
cSSp: contralateral primary somatosensory cortex
cSp5: contralateral spinal trigeminal nucleus
CST: corticospinal tract
CT: corticothalamic
CTB: Cholera toxin subunit B
ET: extratelencephalic
HAR: human accelerated region
HP: hippocampus
HSV: herpes simplex virus
HT: hypothalamus
HPF: hippocampal formation
i-ECT: ipsilateral ectorhinal cortex
ILA: infralimbic area
i-MO: ipsilateral motor cortex
iMOs: ipsilateral secondary motor cortex
IP: intermediate progenitor
IS: intersection/subtraction
IT: intratelencephalic
KI: knock-in
LA: lateral amygdala
MD: mediodorsal nucleus of the thalamus
MO: motor cortex
MOp: Primary motor cortex
MOs: secondary motor cortex
nRG: neurogenic radial glia
OB: olfactory bulb
OLF: olfactory areas
ORBm: Orbital area medial part
ORBl/vl: Orbital area lateral/ventral lateral parts
PAL: pallidum
PAG: periaqueductal gray
PL: prelimbic area
Pn: pons
POm: posteromedial complex of the thalamus
PT: pyramidal tract
PV: parvalbumin
PyN: pyramidal neuron
retroAAV: retrograde adeno-associated virus
RG: radial glia
RT: reticular nucleus of the thalamus
SC: superior colliculus
SCPN: subcerebral projection neurons
Spd: spinal cord
SSp: Primary somatosensory cortex
SSp-bfd: Primary somatosensory barrelfield cortex
SSs: secondary somatosensory cortex
STP: serial two-photon
Str: striatum
SVZ: subventricular zone
TEa: temporal association area
TF: transcription factor
Th/THAL/Thal: thalamus
TM: tamoxifen
TRE: tetracycline-responsive promoter element
tTA: tetracyclin controlled transactivator
VISp: Primary visual cortex
VM: ventromedial nucleus of the thalamus
VPM: ventral posteromedial nucleus of the thalamus
VZ: ventricular zone
ZI: zona incerta

## METHODS

### Generation of New Knockin Mouse Lines

Driver and reporter mouse lines listed in Table 1 were generated using a PCR-based cloning, as described before and below ^28, 60, 78^. All experimental procedures were approved by the Institutional Animal Care and Use Committee (IACUC) of CSHL in accordance with NIH guidelines. Mouse knockin driver lines are deposited at Jackson Laboratory for wide distribution. Knockin mouse lines were generated by inserting a 2A-CreER or 2A-Flp cassette in-frame before the STOP codon of the targeted gene. Targeting vectors were generated using PCR-based cloning approach as described before. Briefly, for each gene of interest, two partially overlapping BAC clones from the RPCI-23&24 library (made from C57BL/b mice) were chosen from the Mouse Genome Browser. 5’ and 3’ homology arms were PCR amplified (2-5kb upstream and downstream, respectively) using the BAC DNA as template and cloned into a building vector to flank the 2A-CreERT2 or 2A-Flp expressing cassette as described ^78^. These targeting vectors were purified, tested for integrity by enzyme restriction and PCR sequencing. Linearized targeting vectors were electroporated into a 129SVj/B6 hybrid ES cell line (v6.5). ES clones were first screened by PCR and then confirmed by Southern blotting using appropriate probes. DIG-labeled Southern probes were generated by PCR, subcloned, and tested on wild-type genomic DNA to verify that they give clear and expected results. Positive v6.5 ES cell clones were used for tetraploid complementation to obtain male heterozygous mice following standard procedures. The F0 males were bred with reporter lines (Supplementary Tables 1, 3, 4) and induced with tamoxifen at the appropriate ages to characterize the resulting genetically targeted recombination patterns.

### Tamoxifen Induction

Tamoxifen (T5648, Sigma) was prepared by dissolving in corn oil (20 mg•ml^-1^), applying a sonication pulse for 60 s, followed by constant rotation overnight at 37°C. Embryonic inductions for most Knockin lines were done in Swiss-Webster background; inductions for *Tis21*-CreER, *Fezf2*-Flp intersection experiments were done in C57Bl6 background. E0.5 was established as noon on the day of vaginal plug and tamoxifen was administered to pregnant mothers by gavage at a dose varying from 2-100 mg•kg^-1^ at the appropriate age. For embryonic harvests (12-24 hour pulse chase experiments), a dose of 2mg•kg^-1^ was administered to pregnant moms via oral gavage. For postnatal induction, a 100-200 mg•kg^-1^ dose was administered by intraperitoneal injection at the appropriate age.

### Immunohistochemistry

Postnatal and adult mice were anesthetized (using Avertin) and intracardially perfused with saline followed by 4% paraformaldehyde (PFA) in 0.1 M PB. Following overnight post-fixation at 4°C, brains were rinsed three times and sectioned 50-75 μm thick with a Leica 1000s vibratome. Embryonic brains were collected in PBS and fixed in 4% paraformaldehyde (PFA) 4h at room temperature, rinsed three times with PBS, dehydrated in 30% sucrose-PBS, frozen in OCT compound and cut by cryostat (Leica, CM3050S) in 20-50μm coronal sections. Early postnatal pups were anaesthetized using cold shock on ice and intracardially perfused with 4% PFA in PBS. Post-fixation was performed similar to older mice. Postnatal mice aged 1-2 months were anesthetized using Avertin and intracardially perfused with saline followed by 4% PFA in PBS; brains were post-fixed in 4% PFA overnight at 4 °C and subsequently rinsed three times, embedded in 3% agarose-PBS and cut 50–100 μm in thickness using a vibrating microtome (Leica, VT100S). Sections were placed in blocking solution containing 10% Normal Goat Serum (NGS) and 0.1% Triton-X100 in PBS1X for 1 hr, then incubated overnight at 4 °C with primary antibodies diluted blocking solution. Sections were rinsed 3 times in PBS and incubated for 1 h at room temperature with corresponding secondary antibodies (1:500, Life Technologies). Sections were washed three times with PBS and incubated with DAPI for 5 min (1:5,000 in PBS, Life Technologies, 33342) to stain nuclei. Sections were dry-mounted on slides using Vectashield (Vector Labs, H1000) or Fluoromount (Sigma, F4680) mounting medium.

To perform molecular characterization of GeneX-CreER mouse lines, we stained 40μm vibratome sections for Cux1 and Ctip2, that were imaged in a Nikon Eclipse 90i fluorescence microscope. Focusing on the somatosensory cortex, we counted tdTomato+ cells in a ∼300μm width column and determined their relative position along the dorso-ventral axis that goes from the ventricular surface (0) to the pia (100%). As a reference, Ctip2+ and Cux1+ regions were plotted as green and blue bars, where the upper limits correspond to the mean relative position of the dorsal-most positive cells, and the lower limits correspond to the mean relative position of the ventral-most positive cells. Gray areas in histograms correspond to SD of those limits. The frequency of tdTomato+ cells along the dorso-ventral axis was plotted in a histogram with a bin width of 5%. Number of cells: *Fezf2*-CreER: 2781 cells, *Tcerg1l*-CreER: 185 cells, *Adcyap1*-CreER: 54 cells, *Tle4*-CreER: 2737 cells, *Lhx2*-CreER: 1380 cells, *PlexinD1*-CreER: 809 cells, *Cux1*-CreER: 2296 cells, *Tbr1*-CreER: 3572 cells, *Tbr2*-CreER TM E16.5: 1273 cells, *Tbr2*-CreER TM E17.5: 1871 cells. For each line we quantified at least 4 sections from 2 embryos. Differences in cell numbers are due to differences in labeling density.

For colocalization determination, we obtained confocal z stacks centered in layer 5 or 6 of the somatosensory cortex, of 320×320×40μm^3^ volumes. For all tdTomato+ cells in the volume we manually determined whether they were also positive for the desired markers by looking in individual z planes. The percentage of positive cells was calculated for each area. Average number of tdTomato+ cells quantified per staining: Fezf2-CreER: 314 cells in layer 5 and 472 in layer 6, Tcerg1l-CreER: 162 cells, Adcyap-CreER: 20 cells, Tle4-CreER: 157 cells in layer 5 and 1081 in layer 6, Lhx2-CreER: 294 cells, PlexinD1-CreER: 468 cells in layers 4 and 5a, Cux1-CreER: 761 cells, Tbr1-CreER: 858 cells, Tbr2-CreER: 1380 cells. For each line we quantified at least 4 sections from at least 2 embryos. Differences in cell numbers are due to differences in labeling density.

### Antibodies

Anti-GFP (1:1000, Aves, GFP-1020); anti-RFP (1:1000, Rockland Pharmaceuticals, 600-401-379); anti-mCherry (1:500, OriGene AB0081-500); anti-mKate2 for Brainbow 3.0 (gift of Dr. Dawen Cai, U Michigan); anti-SATB2 (1:20, Abcam ab51502); anti-CTIP2 (1:100, Abcam 18465); anti-CUX1 (1:100, SantaCruz 13024); anti-LDB2 (1:200, Proteintech 118731-AP); anti-Fog2 (1:500, SantaCruz m-247), anti-LHX2 (1:250, Millipore-Sigma ABE1402) and anti-Tle4 (1:300, Santa Cruz sc-365406) were used.

### Validation of PyN driver lines

ViewRNA tissue Assay (Thermo Fisher Scientific) Fluorescent in situ hybridization (FISH) was carried out per manufacturer’s instructions on genetically identified PyNs expressing H2bGFP nuclear reporter (GeneX-CreER;LSL-H2bGFP) to validate the expression of PyN mRNA within cre-recombinase dependent H2bGFP expressing cells in adult tissue (p24). Antibody validation with cre-recombinase dependent reporter (GeneX-CreER;Ai14) was also used as was available for use in adult tissue. For both FISH and antibody validation experiments, the percentage of total recombinase dependent reporter positive cells co-expressing PyN driver transcript or antibody was quantified.

### Viral Injection and Analysis

#### Stereotaxic viral injection

Adult mice were anesthetized by inhalation of 2% isofluorane delivered with a constant air flow (0.4 L•min^-1^). Ketoprofen (5 mg•kg^-1^) and dexamethasone (0.5 mg•kg^-1^) were administered subcutaneously as preemptive analgesia and to prevent brain edema, respectively, prior to surgery, and lidocaine (2-4 mg•kg^-1^) was applied intra-incisionally. Mice were mounted in a stereotaxic headframe (Kopf Instruments, 940 series or Leica Biosystems, Angle Two). Stereotactic coordinates were identified (Supplementary Table 5). An incision was made over the scalp, a small burr hole drilled in the skull and brain surface exposed. Injections were performed according to the strategies delineated in Supplementary Table 5. A pulled glass pipette tip of 20–30 μm containing the viral suspension was lowered into the brain; a 300-400 nl volume was delivered at a rate of 30 nl•min^-1^ using a Picospritzer (General Valve Corp); the pipette remained in place for 10 min preventing backflow, prior to retraction, after which the incision was closed with 5/0 nylon suture thread (Ethilon Nylon Suture, Ethicon Inc. Germany) or Tissueglue (3M Vetbond), and animals were kept warm on a heating pad until complete recovery.

#### Systemic AAV injection

Foxp2-IRES-Cre mice were injected through the lateral tail vein at 4 weeks of age with 100 μl total volume of AAV9-CAG-DIO-EGFP (UNC Viral Core) diluted in PBS (5×1011 vg/mouse). Three weeks postinjection, mice were transcardially perfused with 0.9% saline, followed by ice-cold 4% PFA in PBS, and processed for STP tomography.

### Virus

Adeno-associated viruses (AAVs) serotype 8, 9, DJ PHP.eB or rAAV2-retro (retroAAV) packaged by commercial vector core facilities (UNC Vector Core, ETH Zurich, Biohippo, Penn, Addgene) were used as listed in Supplementary Table 5. Briefly, for cell-type specific anterograde tracing, we used either Cre- or Flp-dependent or tTA-activated AAVs combined with the appropriate reporter mouse lines (Supplementary Table 7) ^28^, or dual-tTA (Fig. 5 & 6) to express EGFP, EYFP or mRuby2 in labeled axons. retroAAV-Flp was used to infect axons at their terminals^6^ in target brain structures to label PyNs retrogradely according to experiments detailed in Supplementary Table 5.

### Microscopy and Image Analysis

Imaging was performed using Zeiss LSM 780 or 710 confocal microscopes, Nikon Eclipse 90i or Zeiss Axioimager M2 fluorescence microscopes, or whole brain Serial Two-Photon (STP) Tomography (detailed in section below). Imaging from serially mounted sections was performed on a Zeiss LSM 780 or 710 confocal microscope (CSHL St. Giles Advanced Microscopy Center) and Nikon Eclipse 90i fluorescence microscope, using objectives X63 and x5 for embryonic tissue, and x20 for adult tissue, as well as x5 on a Zeiss Axioimager M2 System equipped with MBF Neurolucida Software (MBF). Quantification and image analysis was performed using Image J/FIJI software. Statistics and plotting of graphs were done using GraphPad Prism 7 and Microsoft Excel 2010.

#### Target-specific Cell Depth Measurement

Cell depth analysis for retrogradely labeled projection specific genetically identified PyNs (GeneX-CreER) were obtained using 5x MBF fluorescent widefield images of 65-μm thick coronal sections in MO and SSp-bfd. MO cell depths are presented in microns due to the absence of a defined white matter border in frontal cortical areas and SSp-bfd depth ratio measurements were normalized to the distance from pia to white matter. For each condition we quantified at least 4 sections taken from 2 animals.

### Whole-brain Serial Two Photon Tomography and Image Analysis

Perfused and post-fixed brains from adult mice were embedded in oxidized agarose and imaged with TissueCyte 1000 (Tissuevision) as described ^4, 5^. We used the whole-brain STP tomography pipeline previously described ^4, 5^. Perfused and post-fixed brains from adult mice, prepared as described above, were embedded in 4% oxidized-agarose in 0.05M PB, cross-linked in 0.2% sodium borohydrate solution (in 0.05 M sodium borate buffer, pH 9.0-9.5). The entire brain was imaged in coronal sections with a 20x Olympus XLUMPLFLN20XW lens (NA 1.0) on a TissueCyte 1000 (Tissuevision) with a Chameleon Ultrafast-2 Ti: Sapphire laser (Coherent). EGFP/EYFP or tdTomato signals were excited at 910 nm or 920 nm, respectively. Whole brain image sets were acquired as series of 12 (x) x 16 (y) tiles with 1 μm x 1 μm sampling for 230-270 z sections with a 50-μm z-step size. Images were collected by two PMTs (PMT, Hamamatsu, R3896), for signal and autofluorescent background, using a 560 nm dichroic mirror (Chroma, T560LPXR) and band pass filters (Semrock FF01-680/SP-25). The image tiles were corrected to remove illumination artifacts along the edges and stitched as a grid sequence ^79, 80^. Image processing was completed using ImageJ/FIJI and Adobe/Photoshop software with linear level and nonlinear curve adjustments applied only to entire images.

#### Cell body detection from whole-brain STP data

PyN somata were automatically detected from cell-type specific reporter lines (R26-LSL-GFP or Ai14) by a convolutional network trained as described in ^81^. Detected PyN soma coordinates were overlaid on a mask for cortical depth, as described ^81^.

#### Axon detection from whole-brain STP data

For axon projection mapping, PyN axon signal based on cell-type specific viral expression of EGFP or EYFP was filtered by applying a square root transformation, histogram matching to the original image, and median and Gaussian filtering using Fiji/ImageJ software ^82^ so as to maximize signal detection while minimizing background auto-fluorescence, as described in ^83^. A normalized substraction of the autofluorescent background channel was applied and the resulting thresholded images were converted to binary maps. 3D rendering was performed based on binarized axon projections and surfaces were determined based on the binary images using Imaris software (Bitplane). Projections were quantified as the fraction of pixels in each brain structure relative to each whole projection.

#### Registration of whole-brain STP image datasets

Registration brain-wide datasets to the Allen reference Common Coordinate Framework (CCF) version 3 was performed by 3D affine registration followed by a 3D B-spline registration using Elastix software ^84^, according to established parameters ^84^. For cortical depth and axon projection analysis, we registered the CCFv3 to each dataset so as to report cells detected and pixels from axon segmentation in each brain structure without warping the imaging channel.

### In vitro Electrophysiology

#### Brain slice preparation

Mice (>P30) were anesthetized with isoflurane, decapitated, brains dissected out and rapidly immersed in ice-cold, oxygenated, artificial cerebrospinal fluid (section ACSF: 110 mM choline-Cl, 2.5 mM KCl, 4mM MgSO4, 1mM CaCl2, 1.25 mM NaH2PO4, 26mM NaHCO3, 11mM D-glucose, 10 mM Na ascorbate, 3.1 Na pyruvate, pH 7.35, 300 mOsm) for 1 min. Coronal cortical slices containing somatomotor cortex were sectioned at 300 μm thickness using a vibratome (HM 650 V; Microm) at 1-2 °C and incubated with oxygenated ACSF (working ACSF; 124mM NaCl, 2.5 mM KCl, 2 mM MgSO4, 2 mM CaCl2, 1.25 mM NaH2PO4, 26 mM NaHCO3, 11 mM D-glucose, pH 7.35, 300mOsm) at 34 °C for 30 min, and subsequently transferred to ACSF at room temperature (25 °C) for >30 min before use. Whole cell patch recordings were directed to the somatosensory and motor cortex, the subcortical whiter matter and corpus callosum served as primary landmarks according to the atlas (Paxinos and Watson Mouse Brain in Stereotaxic Coordinates, 3rd edition).

#### Patch clamp recording in brain slice

Patch pipettes were pulled from borosilicate glass capillaries with filament (1.2 mm outer diameter and 0.69 inner diameter; Warner Instruments) with a resistance of 3-6 MΩ. The pipette recording solution consisted of 130 mM potassium gluconate, 15 mM KCl, 10 mM sodium phosphocreatine, 10 mM Hepes, 4 mM ATP·Mg, 0.3 mM GTP, and 0.3 mM EGTA (pH 7.3 adjusted with KOH, 300 mOsm). Dual or triple whole cell recordings from tdTomato+ and EGFP+ PyNs were made with Axopatch 700B amplifiers (Molecular Devices, Union City, CA) using an upright microscope (Olympus, Bx51) equipped with infrared-differential interference contrast optics (IR-DIC) and fluorescence excitation source. Both IR-DIC and fluorescence images were captured with a digital camera (Microfire, Optronics, CA). All recordings were performed at 33–34 °C with the chamber perfused with oxygenated working ACSF.

Recordings were made with two MultiClamp 700B amplifiers (Molecular Devices). The membrane potential was maintained at −75mV in the voltage clamping mode and zero holding current in the current clamping mode, without the correction of junction potential. Signals were recorded and filtered at 2 kHz, digitalized at 20 kHz (DIGIDATA 1322A, Molecular Devices) and further analyzed using the pClamp 10.3 software (Molecular Devices) for intrinsic properties.

## DATA AVAILABILITY

Raw and stitched whole-brain STP imaging data is available from the BICCN Brain Image Library (BIL) (http://www.brainimagelibrary.org/download.html) at the Pittsburgh Supercomputing Center. Anterograde projection datasets can be visualized on the Mouse Brain architecture website (http://brainarchitecture.org/cell-type/projection) as detailed in Supplementary Tables 5, 6.

## ACKNOWLEDGMENTS

We are grateful to Dr. Richard Palmiter for providing the *FoxP2-IRES-Cre* mouse line and to Dr. Yutaka Yoshida for providing the *Sema3E-CreER* mouse line. We thank Dr. Leyi Li at CSHL for help with generation of knock-in mice, Dr. Sean M. Kelly for contribution to generating and characterizing the *Tis21-2A-CreER* mouse line, Dr. Yongsoo Kim for providing the digital flatmap of the mouse cortex, Dr. Francesco Boato for help in mouse breeding and brain tissue preparation, Dr. Kathleen Kelly for project management, H. Kondo for providing anatomical datasets, S. Srivas for help with Tbr1 driver data, B. Huo, X. Li, F. Xu for help with data sharing. We thank CSHL Laboratory Animal Resources for mouse husbandry. This work was supported in part by the NIH grants 5R01MH101268-05 and 5U19MH114821-03 to Z.J.H. and P.A., 1S10OD021759-01 to Z.J.H., the CSHL Robertson Neuroscience Fund to Z.J.H. P.O. is supported by NIH U01 MH114824-01. D.H. was supported by Human Frontier Science Program long term fellowship LT000075/2014-L and NARSAD Young Investigator grant #26327. R.R. was supported by NRSA F31 Predoctoral Fellowship 5F31MH114529-03. J.T. and J.M.L. were supported by NRSA F30 Medical Scientist Predoctoral Fellowships 5F30MH097425-03 and 5F30MH108333.

## AUTHOR CONTRIBUTIONS

Z.J.H and P.A. conceived the study and obtained funding. Z.J.H. coordinated the study.

Z.J.H., P.A., K.M., D.H. designed the experiments. M.H., P.W and E.G. designed and generated all new knock-in mouse lines. K.M., X.A., D.H., R.P., J.H., D.D., J.T., J.M.L., R.R., W.G., E.G., P.W., M.H., J.A.H., and E.S. conducted mouse breeding, anatomy, immunohistochemistry, imaging, and quantification.

K.M., W.G., X.A., G.K., J.H. and R.P. conducted virus injection experiments.

J.L. performed the patch clamp recording in brain slice. K.M., J.H., R.P., X.A., W.G., A.N., P.M. and P.O. performed whole-brain Serial Two-Photon Tomography and analysis of cell distribution and axon projection patterns. Z.J.H. wrote the manuscript with contributions and edits from P.A., K.M., D.H., W.G., M.H., J.L., X.A.

## REFERENCES

1. Harris, K. D. & Shepherd, G. M. The neocortical circuit: themes and variations. Nat Neurosci 18, 170–181, doi:10.1038/nn.3917 (2015).

2. Huang, Z. J. Toward a genetic dissection of cortical circuits in the mouse. Neuron 83, 1284–1302, doi:10.1016/j.neuron.2014.08.041 (2014).

3. Lodato, S. & Arlotta, P. Generating neuronal diversity in the mammalian cerebral cortex. Annu Rev Cell Dev Biol 31, 699–720, doi:10.1146/annurev-cellbio-100814-125353 (2015).

4. Gerfen, C. R., Economo, M. N. & Chandrashekar, J. Long distance projections of cortical pyramidal neurons. J Neurosci Res 96, 1467–1475, doi:10.1002/jnr.23978 (2018).

5. Kawaguchi, Y. Pyramidal Cell Subtypes and Their Synaptic Connections in Layer 5 of Rat Frontal Cortex. Cereb Cortex 27, 5755–5771, doi:10.1093/cercor/bhx252 (2017).

6. Tasic, B. et al. Shared and distinct transcriptomic cell types across neocortical areas. Nature 563, 72–78, doi:10.1038/s41586-018-0654-5 (2018).

7. Rakic, P. The radial edifice of cortical architecture: from neuronal silhouettes to genetic engineering. Brain Res Rev 55, 204–219, doi:S0165-0173(07)00035-5 [pii] 10.1016/j.brainresrev.2007.02.010 (2007).

8. Kriegstein, A., Noctor, S. & Martinez-Cerdeno, V. Patterns of neural stem and progenitor cell division may underlie evolutionary cortical expansion. Nat Rev Neurosci 7, 883–890, doi:nrn2008 [pii] 10.1038/nrn2008 (2006).

9. Telley, L. et al. Temporal patterning of apical progenitors and their daughter neurons in the developing neocortex. Science 364, doi:10.1126/science.aav2522 (2019).

10. Greig, L. C., Woodworth, M. B., Galazo, M. J., Padmanabhan, H. & Macklis, J. D. Molecular logic of neocortical projection neuron specification, development and diversity. Nat Rev Neurosci 14, 755–769, doi:10.1038/nrn3586 (2013).

11. Eckler, M. J. et al. Cux2-positive radial glial cells generate diverse subtypes of neocortical projection neurons and macroglia. Neuron 86, 1100–1108, doi:10.1016/j.neuron.2015.04.020 (2015).

12. Franco, S. J. et al. Fate-restricted neural progenitors in the mammalian cerebral cortex. Science 337, 746–749, doi:337/6095/746 [pii]10.1126/science.1223616.

13. Gil-Sanz, C. et al. Lineage Tracing Using Cux2-Cre and Cux2-CreERT2 Mice. Neuron 86, 1091–1099, doi:10.1016/j.neuron.2015.04.019 (2015).

14. Guo, C. et al. Fezf2 expression identifies a multipotent progenitor for neocortical projection neurons, astrocytes, and oligodendrocytes. Neuron 80, 1167–1174, doi:10.1016/j.neuron.2013.09.037 (2013).

15. Gerfen, C. R., Paletzki, R. & Heintz, N. GENSAT BAC cre-recombinase driver lines to study the functional organization of cerebral cortical and basal ganglia circuits. Neuron 80, 1368–1383, doi:10.1016/j.neuron.2013.10.016 (2013).

16. Harris, J. A. et al. Anatomical characterization of Cre driver mice for neural circuit mapping and manipulation. Front Neural Circuits 8, 76, doi:10.3389/fncir.2014.00076 (2014).

17. Chou, S. J. & Tole, S. Lhx2, an evolutionarily conserved, multifunctional regulator of forebrain development. Brain Res 1705, 1–14, doi:10.1016/j.brainres.2018.02.046 (2019).

18. Woodworth, M. B., Greig, L. C., Kriegstein, A. R. & Macklis, J. D. SnapShot: cortical development. Cell 151, 918–918 e911, doi:10.1016/j.cell.2012.10.004 (2012).

19. Bulchand, S., Grove, E. A., Porter, F. D. & Tole, S. LIM-homeodomain gene Lhx2 regulates the formation of the cortical hem. Mech Dev 100, 165–175, doi:10.1016/s0925-4773(00)00515-3 (2001).

20. Mangale, V. S. et al. Lhx2 selector activity specifies cortical identity and suppresses hippocampal organizer fate. Science 319, 304–309, doi:10.1126/science.1151695 (2008).

21. Monuki, E. S., Porter, F. D. & Walsh, C. A. Patterning of the dorsal telencephalon and cerebral cortex by a roof plate-Lhx2 pathway. Neuron 32, 591–604, doi:10.1016/s0896-6273(01)00504-9 (2001).

22. Shetty, A. S. et al. Lhx2 regulates a cortex-specific mechanism for barrel formation. Proc Natl Acad Sci U S A 110, E4913–4921, doi:10.1073/pnas.1311158110 (2013).

23. Chen, J. G., Rasin, M. R., Kwan, K. Y. & Sestan, N. Zfp312 is required for subcortical axonal projections and dendritic morphology of deep-layer pyramidal neurons of the cerebral cortex. Proc Natl Acad Sci U S A 102, 17792–17797, doi:0509032102 [pii] 10.1073/pnas.0509032102 (2005).

24. Molyneaux, B. J., Arlotta, P., Hirata, T., Hibi, M. & Macklis, J. D. Fezl is required for the birth and specification of corticospinal motor neurons. Neuron 47, 817–831, doi:S0896-6273(05)00732-4 [pii] 10.1016/j.neuron.2005.08.030 (2005).

25. Muralidharan, B. et al. LHX2 Interacts with the NuRD Complex and Regulates Cortical Neuron Subtype Determinants Fezf2 and Sox11. J Neurosci 37, 194–203, doi:10.1523/JNEUROSCI.2836-16.2016 (2017).

26. Chou, S. J., Perez-Garcia, C. G., Kroll, T. T. & O’Leary, D. D. Lhx2 specifies regional fate in Emx1 lineage of telencephalic progenitors generating cerebral cortex. Nat Neurosci 12, 1381–1389, doi:10.1038/nn.2427 (2009).

27. Godbole, G. et al. Hierarchical genetic interactions between FOXG1 and LHX2 regulate the formation of the cortical hem in the developing telencephalon. Development 145, doi:10.1242/dev.154583 (2018).

28. He, M. et al. Strategies and Tools for Combinatorial Targeting of GABAergic Neurons in Mouse Cerebral Cortex. Neuron 91, 1228–1243, doi:10.1016/j.neuron.2016.08.021 (2016).

29. Florio, M. & Huttner, W. B. Neural progenitors, neurogenesis and the evolution of the neocortex. Development 141, 2182–2194, doi:10.1242/dev.090571 (2014).

30. Haubensak, W., Attardo, A., Denk, W. & Huttner, W. B. Neurons arise in the basal neuroepithelium of the early mammalian telencephalon: a major site of neurogenesis. Proc Natl Acad Sci U S A 101, 3196–3201, doi:10.1073/pnas.0308600100 (2004).

31. Martinez-Cerdeno, V. et al. Evolutionary origin of Tbr2-expressing precursor cells and the subventricular zone in the developing cortex. J Comp Neurol 524, 433–447, doi:10.1002/cne.23879 (2016).

32. Martinez-Cerdeno, V., Noctor, S. C. & Kriegstein, A. R. The role of intermediate progenitor cells in the evolutionary expansion of the cerebral cortex. Cereb Cortex 16 Suppl 1, i152–161, doi:10.1093/cercor/bhk017 (2006).

33. Sun, T. & Hevner, R. F. Growth and folding of the mammalian cerebral cortex: from molecules to malformations. Nat Rev Neurosci 15, 217–232, doi:10.1038/nrn3707 (2014).

34. Kowalczyk, T. et al. Intermediate neuronal progenitors (basal progenitors) produce pyramidal-projection neurons for all layers of cerebral cortex. Cereb Cortex 19, 2439–2450, doi:10.1093/cercor/bhn260 (2009).

35. Mihalas, A. B. & Hevner, R. F. Control of Neuronal Development by T-Box Genes in the Brain. Curr Top Dev Biol 122, 279–312, doi:10.1016/bs.ctdb.2016.08.001 (2017).

36. Vasistha, N. A. et al. Cortical and Clonal Contribution of Tbr2 Expressing Progenitors in the Developing Mouse Brain. Cereb Cortex 25, 3290–3302, doi:10.1093/cercor/bhu125 (2015).

37. Englund, C. et al. Pax6, Tbr2, and Tbr1 are expressed sequentially by radial glia, intermediate progenitor cells, and postmitotic neurons in developing neocortex. J Neurosci 25, 247–251, doi:10.1523/JNEUROSCI.2899-04.2005 (2005).

38. Kawaguchi, A. et al. Single-cell gene profiling defines differential progenitor subclasses in mammalian neurogenesis. Development 135, 3113–3124, doi:10.1242/dev.022616 (2008).

39. Noctor, S. C., Martinez-Cerdeno, V., Ivic, L. & Kriegstein, A. R. Cortical neurons arise in symmetric and asymmetric division zones and migrate through specific phases. Nature neuroscience 7, 136–144, doi:10.1038/nn1172 nn1172 [pii] (2004).

40. Arlotta, P. et al. Neuronal subtype-specific genes that control corticospinal motor neuron development in vivo. Neuron 45, 207–221, doi:10.1016/j.neuron.2004.12.036 (2005).

41. Arlotta, P., Molyneaux, B. J., Jabaudon, D., Yoshida, Y. & Macklis, J. D. Ctip2 controls the differentiation of medium spiny neurons and the establishment of the cellular architecture of the striatum. J Neurosci 28, 622–632, doi:28/3/622 [pii] 10.1523/JNEUROSCI.2986-07.2008 (2008).

42. Fame, R. M., MacDonald, J. L. & Macklis, J. D. Development, specification, and diversity of callosal projection neurons. Trends Neurosci 34, 41–50, doi:S0166-2236(10)00147-5 [pii] 10.1016/j.tins.2010.10.002 (2011).

43. Hintiryan, H. et al. The mouse cortico-striatal projectome. Nat Neurosci 19, 1100–1114, doi:10.1038/nn.4332 (2016).

44. Hooks, B. M. et al. Topographic precision in sensory and motor corticostriatal projections varies across cell type and cortical area. Nat Commun 9, 3549, doi:10.1038/s41467-018-05780-7 (2018).

45. Zingg, B. et al. Neural networks of the mouse neocortex. Cell 156, 1096–1111, doi:10.1016/j.cell.2014.02.023 (2014).

46. Iwamoto, M., Bjorklund, T., Lundberg, C., Kirik, D. & Wandless, T. J. A general chemical method to regulate protein stability in the mammalian central nervous system. Chem Biol 17, 981–988, doi:S1074-5521(10)00305-4 [pii] 10.1016/j.chembiol.2010.07.009 (2010).

47. Nieto, M. et al. Expression of Cux-1 and Cux-2 in the subventricular zone and upper layers II-IV of the cerebral cortex. J Comp Neurol 479, 168–180, doi:10.1002/cne.20322 (2004).

48. Weiss, L. A. & Nieto, M. The crux of Cux genes in neuronal function and plasticity. Brain Res 1705, 32–42, doi:10.1016/j.brainres.2018.02.044 (2019).

49. Cubelos, B. et al. Cux-2 controls the proliferation of neuronal intermediate precursors of the cortical subventricular zone. Cereb Cortex 18, 1758–1770, doi:10.1093/cercor/bhm199 (2008).

50. Cubelos, B. et al. Cux1 and Cux2 regulate dendritic branching, spine morphology, and synapses of the upper layer neurons of the cortex. Neuron 66, 523–535, doi:10.1016/j.neuron.2010.04.038 (2010).

51. Rodriguez-Tornos, F. M. et al. Cux1 Enables Interhemispheric Connections of Layer II/III Neurons by Regulating Kv1-Dependent Firing. Neuron 89, 494–506, doi:10.1016/j.neuron.2015.12.020 (2016).

52. Molyneaux, B. J. et al. Novel subtype-specific genes identify distinct subpopulations of callosal projection neurons. J Neurosci 29, 12343–12354, doi:29/39/12343 [pii] 10.1523/JNEUROSCI.6108-08.2009 (2009).

53. Chauvet, S. et al. Gating of Sema3E/PlexinD1 signaling by neuropilin-1 switches axonal repulsion to attraction during brain development. Neuron 56, 807–822, doi:10.1016/j.neuron.2007.10.019 (2007).

54. Ding, J. B., Oh, W. J., Sabatini, B. L. & Gu, C. Semaphorin 3E-Plexin-D1 signaling controls pathway-specific synapse formation in the striatum. Nat Neurosci 15, 215–223, doi:10.1038/nn.3003 (2011).

55. Oh, W. J. & Gu, C. The role and mechanism-of-action of Sema3E and Plexin-D1 in vascular and neural development. Semin Cell Dev Biol 24, 156–162, doi:10.1016/j.semcdb.2012.12.001 (2013).

56. Leone, D. P., Srinivasan, K., Chen, B., Alcamo, E. & McConnell, S. K. The determination of projection neuron identity in the developing cerebral cortex. Curr Opin Neurobiol 18, 28–35, doi:10.1016/j.conb.2008.05.006 (2008).

57. Lodato, S. et al. Gene co-regulation by Fezf2 selects neurotransmitter identity and connectivity of corticospinal neurons. Nat Neurosci 17, 1046–1054, doi:10.1038/nn.3757 (2014).

58. Di Bella, D. J. et al. Molecular Logic of Cellular Diversification in the Mammalian Cerebral Cortex. bioRxiv (2020).

59. Watakabe, A., Ohsawa, S., Hashikawa, T. & Yamamori, T. Binding and complementary expression patterns of semaphorin 3E and plexin D1 in the mature neocortices of mice and monkeys. J Comp Neurol 499, 258–273, doi:10.1002/cne.21106 (2006).

60. Pecho-Vrieseling, E., Sigrist, M., Yoshida, Y., Jessell, T. M. & Arber, S. Specificity of sensory-motor connections encoded by Sema3e-Plxnd1 recognition. Nature 459, 842–846, doi:10.1038/nature08000 (2009).

61. Han, W. et al. TBR1 directly represses Fezf2 to control the laminar origin and development of the corticospinal tract. Proc Natl Acad Sci U S A 108, 3041–3046, doi:10.1073/pnas.1016723108 (2011).

62. Galazo, M. J., Emsley, J. G. & Macklis, J. D. Corticothalamic Projection Neuron Development beyond Subtype Specification: Fog2 and Intersectional Controls Regulate Intraclass Neuronal Diversity. Neuron 91, 90–106, doi:10.1016/j.neuron.2016.05.024 (2016).

63. Molyneaux, B. J. et al. DeCoN: genome-wide analysis of in vivo transcriptional dynamics during pyramidal neuron fate selection in neocortex. Neuron 85, 275–288, doi:10.1016/j.neuron.2014.12.024 (2015).

64. Chang, M., Suzuki, N. & Kawai, H. D. Laminar specific gene expression reveals differences in postnatal laminar maturation in mouse auditory, visual, and somatosensory cortex. J Comp Neurol 526, 2257–2284, doi:10.1002/cne.24481 (2018).

65. Kast, R. J., Lanjewar, A. L., Smith, C. D. & Levitt, P. FOXP2 exhibits projection neuron class specific expression, but is not required for multiple aspects of cortical histogenesis. Elife 8, doi:10.7554/eLife.42012 (2019).

66. Rousso, D. L. et al. Two Pairs of ON and OFF Retinal Ganglion Cells Are Defined by Intersectional Patterns of Transcription Factor Expression. Cell Rep 15, 1930–1944, doi:10.1016/j.celrep.2016.04.069 (2016).

67. Co, M., Hickey, S. L., Kulkarni, A., Harper, M. & Konopka, G. Cortical Foxp2 Supports Behavioral Flexibility and Developmental Dopamine D1 Receptor Expression. Cereb Cortex 30, 1855–1870, doi:10.1093/cercor/bhz209 (2020).

68. Economo, M. N. et al. Distinct descending motor cortex pathways and their roles in movement. Nature 563, 79–84, doi:10.1038/s41586-018-0642-9 (2018).

69. Daigle, T. L. et al. A Suite of Transgenic Driver and Reporter Mouse Lines with Enhanced Brain-Cell-Type Targeting and Functionality. Cell 174, 465–480 e422, doi:10.1016/j.cell.2018.06.035 (2018).

70. Laboulaye, M. A., Duan, X., Qiao, M., Whitney, I. E. & Sanes, J. R. Mapping Transgene Insertion Sites Reveals Complex Interactions Between Mouse Transgenes and Neighboring Endogenous Genes. Front Mol Neurosci 11, 385, doi:10.3389/fnmol.2018.00385 (2018).

71. Winnubst, J. et al. Reconstruction of 1,000 Projection Neurons Reveals New Cell Types and Organization of Long-Range Connectivity in the Mouse Brain. Cell 179, 268–281 e213, doi:10.1016/j.cell.2019.07.042 (2019).

72. Peng, H. et al. Brain-wide single neuron reconstruction reveals morphological diversity in molecularly defined striatal, thalamic, cortical and claustral neuron types. BioRxiv (2020).

73. Wang, X. et al. Genetic Single Neuron Anatomy Reveals Fine Granularity of Cortical Axo-Axonic Cells. Cell Rep 26, 3145–3159 e3145, doi:10.1016/j.celrep.2019.02.040 (2019).

74. Arlotta, P. & Hobert, O. Homeotic Transformations of Neuronal Cell Identities. Trends Neurosci 38, 751–762, doi:10.1016/j.tins.2015.10.005 (2015).

75. Hodge, R. D. et al. Conserved cell types with divergent features in human versus mouse cortex. Nature 573, 61–68, doi:10.1038/s41586-019-1506-7 (2019).

76. Doan, R. N. et al. Mutations in Human Accelerated Regions Disrupt Cognition and Social Behavior. Cell 167, 341–354 e312, doi:10.1016/j.cell.2016.08.071 (2016).

77. Blankvoort, S., Witter, M. P., Noonan, J., Cotney, J. & Kentros, C. Marked Diversity of Unique Cortical Enhancers Enables Neuron-Specific Tools by Enhancer-Driven Gene Expression. Curr Biol 28, 2103–2114 e2105, doi:10.1016/j.cub.2018.05.015 (2018).

78. Taniguchi, H. et al. A resource of Cre driver lines for genetic targeting of GABAergic neurons in cerebral cortex. Neuron 71, 995–1013, doi:10.1016/j.neuron.2011.07.026 (2011).

79. Preibisch, S., Saalfeld, S. & Tomancak, P. Globally optimal stitching of tiled 3D microscopic image acquisitions. Bioinformatics 25, 1463–1465, doi:10.1093/bioinformatics/btp184 (2009).

80. Ragan, T. et al. Serial two-photon tomography for automated ex vivo mouse brain imaging. Nat Methods 9, 255–258, doi:10.1038/nmeth.1854 (2012).

81. Kim, Y. et al. Brain-wide Maps Reveal Stereotyped Cell-Type-Based Cortical Architecture and Subcortical Sexual Dimorphism. Cell 171, 456–469 e422, doi:10.1016/j.cell.2017.09.020 (2017).

82. Schindelin, J. et al. Fiji: an open-source platform for biological-image analysis. Nat Methods 9, 676–682, doi:10.1038/nmeth.2019 (2012).

83. Mandelbaum, G. et al. Distinct Cortical-Thalamic-Striatal Circuits through the Parafascicular Nucleus. Neuron 102, 636–652 e637, doi:10.1016/j.neuron.2019.02.035 (2019).

84. Klein, S., Staring, M., Murphy, K., Viergever, M. A. & Pluim, J. P. elastix: a toolbox for intensity-based medical image registration. IEEE Trans Med Imaging 29, 196–205, doi:10.1109/TMI.2009.2035616 (2010).

85. Kinare, V. et al. An evolutionarily conserved Lhx2-Ldb1 interaction regulates the acquisition of hippocampal cell fate and regional identity. Development 147, doi:10.1242/dev.187856 (2020).

86. Eckler, M. J. et al. Multiple conserved regulatory domains promote Fezf2 expression in the developing cerebral cortex. Neural Dev 9, 6, doi:10.1186/1749-8104-9-6 (2014).

